# Cell-Intrinsic and Extrinsic Effects of Galectin-1 and Galectin-3 in B-Cell Precursor Acute Lymphoblastic Leukemia

**DOI:** 10.1101/2021.09.22.461145

**Authors:** Fei Fei, Eun Ji Joo, Lu Yang, John Groffen, Nora Heisterkamp

## Abstract

Acute lymphoblastic leukemias arising from the malignant transformation of B-cell precursors (BCP-ALLs) are protected against chemotherapy by both intrinsic factors as well as by interactions with bone marrow stromal cells. Galectin-1 and Galectin-3 have overlapping expression patterns and potentially common functions in these cells. We used Galectin-1 and Galectin-3 double null mutant murine BCP-ALL cells to examine the effect of loss endogenous Galectins. We also tested the effect of dual Galectin inhibition by use of plant-derived natural compounds GM-CT-01 and GR-MD-02 in human BCP-ALL cells in co-culture with stroma. Transformed wild type and Galectin-1 x Galectin-3 double knockout BCP-ALL cells were similar in immunophenotype and active intracellular signaling. However, compared to wild type cells, they showed impaired migration, significantly reduced proliferation and increased sensitivity to drug treatment. GM-CT-01 and GR-MD-02 attenuated intracellular signal transduction and sensitized human BCP-ALL cells to chemotherapy including vincristine and the targeted tyrosine kinase inhibitor nilotinib. Our data show endogenous and extracellular Galectins contribute positively to the growth and survival of BCP-ALL cells. Strategies that would efficiently ablate these two Galectins at both locations would decrease microenvironmental protection and reduce BCP-ALL persistence in the protective bone marrow niche after chemotherapy.

## 1. Introduction

B-cell precursor acute lymphoblastic leukemias (BCP-ALLs) constitute a group of developmentally arrested, immature B-lineage precursors, which can be categorized into numerous subgroups based on genetic abnormalities (1–4). Ph-chromosome positive ALL (Ph-positive ALL) is one major poor-prognosis subcategory of ALL, characterized by the presence of a t(9;22) translocation which fuses two genes, *BCR* and *ABL*, at the breakpoints (5). The chimeric Bcr/Abl protein that is produced as a result has deregulated tyrosine kinase activity which can be inhibited by targeted small molecule inhibitors such as nilotinib (6). This type of leukemia can be studied in the murine system making use of *BCR/ABL* transgenic mice or by generating BCP-ALL from mouse bone marrow by transduction with a retrovirus encoding human Bcr/Abl (7, 8). Other BCP-ALLs involve transcription factors, such as the fusion of ETV6 and Runx1 (9) or recombination of different genes with the MLL gene (10).

All BCP-ALLs have a common feature: their anatomical site of origin, the bone marrow microenvironment, which is also the most frequent location of relapse during or after chemotherapy (11). In fact, drug resistance development of cancer cells is actively promoted by their microenvironment (12, 13). We and others (14–19) model BCP-ALL drug resistance development *ex vivo*, because BCP-ALL cells from patients can proliferate if supported by bone marrow stromal cells. Rellick *et al* (20) recently reported that cells in such co-cultures have a gene expression signature that mimics that of cells remaining after induction therapy designated minimal residual disease (MRD). Indeed, chemotherapy treatment of co-cultures of primary or PDX-derived human BCP-ALL cells with OP9, a murine bone marrow stromal cell line, also allows the persistence of drug-insensitive cells [e.g., (14, 17, 19, 21–24)]. OP9 cells are a major source of SDF1α, which is the main chemokine attracting leukemia cells and promoting their adhesion in the bone marrow (25, 26). The migration towards and adhesion to bone marrow stroma is one mechanism that contributes to chemoprotection provided by stromal cells to BCP-ALL (27, 28) but overall, the mechanisms through which such bone marrow stromal cells promote BCP-ALL drug resistance are incompletely understood.

Although soluble factors are also important, the close association between the BCP-ALL and stromal cells is needed for optimal protection against chemotherapy. There is increasing evidence that both Galectin-1 and Galectin-3 are involved in the protective cross-communication between leukemia cells and the microenvironment. Our data identified Galectin-3, a carbohydrate-binding protein that cross-links polyLacNAc-modified glycoproteins on the cell surface (29, 30), as promoting migration and adhesion of BCP-ALL cells to stromal cells, which are a source of secreted Galectin-3. We showed that drug-treated BCP-ALL cells also synthesize Galectin-3 endogenously (22) but that stromal-produced Galectin-3 is a major source of protective Galectin-3 (24).

Juszczynski *et al* previously reported that only BCP-ALL cells containing MLL rearrangements express Galectin-1 endogenously (31). Our own studies determined that expression of Galectin-1 is more ubiquitous and is also present in other subcategories of BCP-ALL including Ph-positive ALL (23). Interestingly, Galectin-1 was shown to be a ligand for the pre-B-cell receptor in normal precursor B cells (32). In fact, bone marrow stromal cells that secrete Galectin-1 may represent a specific niche in the bone marrow for normal BCP development (33). However, stromal cells in general produce both Galectin-1 and Galectin-3: examples include equine bone marrow stromal cells (34), and OP9 murine bone marrow stromal cells which produce exosomes containing both Galectins (22). Thus, the Galectin-1 and Galectin-3 found associated with BCP-ALL is complicated in terms of origin: the leukemia cells can express Galectin-1 and Galectin-3 endogenously, but stromal cells that protect the BCP-ALL cells synthesize and secrete both Galectins as well.

Here, we addressed the combined importance of endogenously expressed Galectin-1 and Galectin-3 in BCP-ALL cells. We generated BCP-ALL cells from *Lgals 1 x Lgals3* double null mutant mouse bone marrow precursor B cells by transduction with the Bcr/Abl oncogene. In addition, we sought to inhibit both extracellular Galectin-1 and Galectin-3 making use of function-inhibitory compounds: as reviewed (35), compounds that have inhibitory activity against both Galectin-1 and Galectin-3 include the disaccharide small molecule TD139 (36) and relatively high molecular weight (50–60 kDa) carbohydrates derived from natural plant glycans such as GM-CT-01 and GR-MD-02 (37–40). Since the *in vitro* K_d_ of both compounds (GR-MD-02: Galectin-1 10 μM, Galectin-3 2.9 μM; GR-CT-01: 8 μM and 2.8 μM) was reported to be comparable (35), they should inhibit both extracellular Galectin-1 and Galectin-3. These compounds progressed to testing in clinical trials and therefore we investigated their effect on BCP-ALL cells when these are grown with stromal support as a source of extracellular Galectin-1 and Galectin-3.

## 2. Results

### 2.1. Expression of Galectin-1 and Galectin-3 in BCP-ALL

We investigated if Galectin-1 and Galectin-3 mRNAs are simultaneously expressed in Ph-positive BCP-ALL cells by meta-analysis of human gene array expression data sets (41, 42) for transcripts of these genes. The majority of samples in two gene array data sets, representing a total of 33 primary cases of Ph-positive ALL, simultaneously expressed both Galectin-3 and Galectin-1 mRNAs (not shown). Interestingly, one data set showed a positive correlation (R^2^=0.46 & p<0.01) between expression of these two genes, suggesting that in some cases their expression may be linked (Figure 1a). In the tel-AML1 subcategory of BCP-ALL, expression was also positively correlated (R^2^=0.26 & p<0.0001, R^2^=0.16 & p<0.01) (Figure 1b) but a similar analysis of MLL-rearranged BCP-ALLs showed no correlation (data not shown), indicating that this may be subtype-dependent.

**Figure 1.**
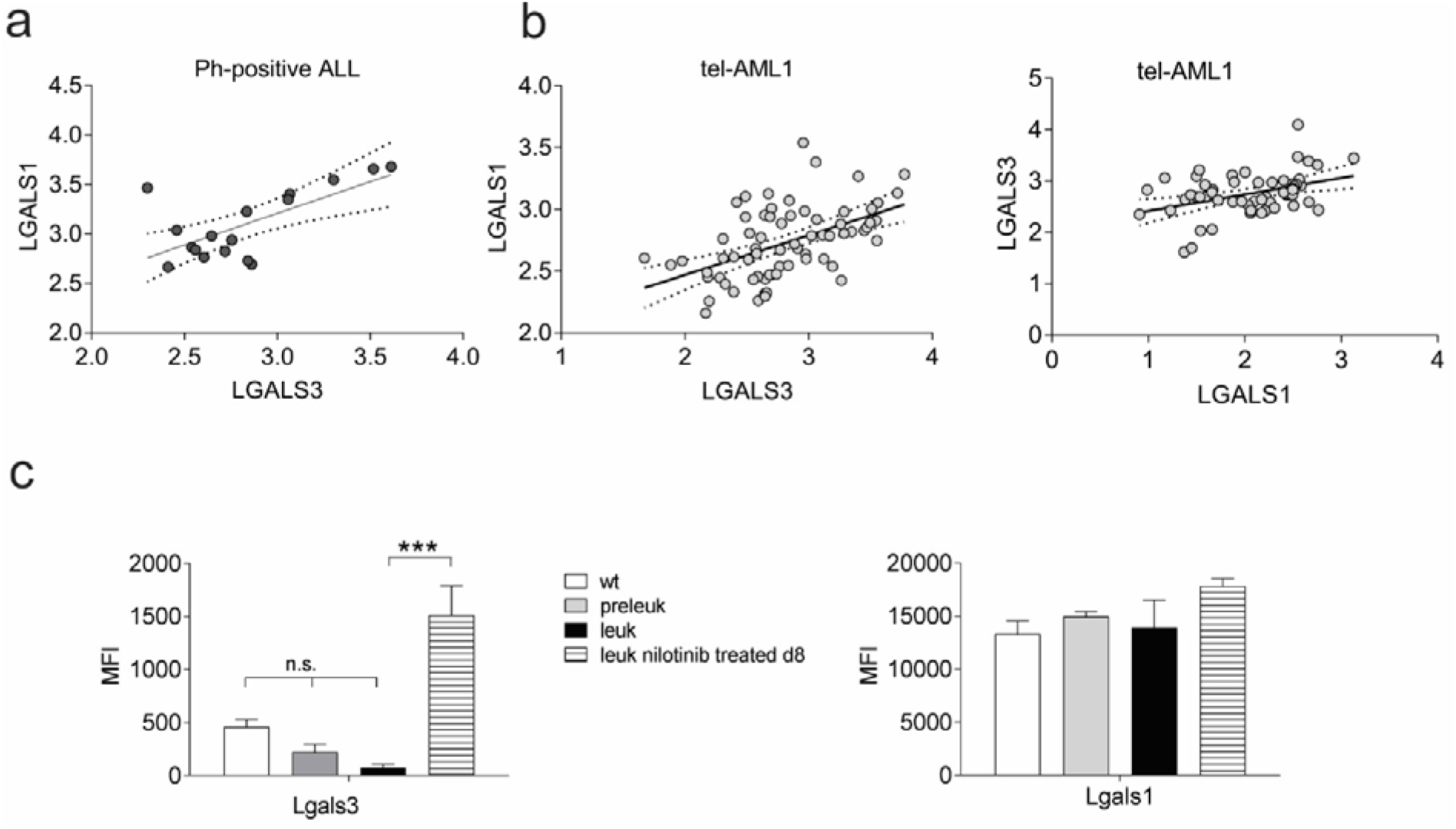
Gene expression of Galectin-1 and Galectin-3 in Bcr-Abl expressing ALL cells. Positive correlation between log10 Galectin-1 and Galectin-3 mRNA expression in (**a**) human Ph-positive ALL samples and (**b**) tel-AML1 ALL samples in two different arrays. Dotted lines indicate 95% confidence range of best fit line. (**c**) Meta-analysis of GSE110104 for Galectin-3 (*Lgals3*) and Galectin-1 (*Lgals1*) gene expression in mouse B-cell precursor cells, flow sorted from bone marrows of control wild type (wt) mice and pre-leukemic, fully leukemic (leuk), and leukemic, 75 mg/kg/d d8 nilotinib-treated *BCR/ABL* P190 transgenic mice. Each set consists of three biological replicates. ***p<0.001. One-way ANOVA, Tukey’s multiple comparisons test. Right panel, no significant differences. MFI, mean fluorescent intensity.

Mouse models for Ph-positive ALL include transgenic mice that develop ALL within around 3 months after birth (7). We analyzed a microarray gene expression data set of such samples (GSE110104) consisting of flow-sorted bone marrow cells from matched wild type, preleukemic, fully leukemia and fully leukemic mice treated for 8 days with nilotinib, a targeted Bcr/Abl kinase inhibitor (43, 44). Whereas the absolute values for Galectin-1 expression were higher than those for Galectin-3 (compare MFI values right and left panel Figure 1c), both non-leukemic wild type and leukemic BCP samples expressed both Galectins. Expression levels of Galectin-1 were comparable in the samples. Consistent with our previous results in human ALL cells treated with drugs (22), the expression of Galectin-3 was significantly induced in drug-treated BCP-ALL cells which we isolated from mice that had been treated for 8 days with 75 mg/kg nilotinib.

### 2.2. Galectin-1 x Galectin-3 double null mutant BCP-ALL cells have decreased proliferation and survival

To investigate the effect of simultaneous loss of Galectin-3 and Galectin-1 function on BCP-ALL cells, we transformed primary B-lineage murine bone marrow cells with the P190 Bcr/Abl oncogene to generate an aggressive, well-proliferating B-cell precursor ALL that does not depend on stromal support. Early passage immunophenotyping of two independently transduced sets of wild type (wt) and *Lgals1 x Lgals3 -/-* (hereafter referred to as dKO) BCP-ALL cells showed they were similar in that they lacked surface IgM, but were positive for CD43, B220 and AA4.1 (Figure 2a). Samples were heterogeneous for expression of the BAFF-R and CD24 (Figure S1 panel a). Western blotting of early passage cells grown without stroma confirmed lack of Galectin-1 (Figure 2b). As expected, based on expression of the Bcr/Abl tyrosine kinase, these cells contained many tyrosine phosphorylated proteins (Figure 2b, pY20 Western blot panel) as a consequence of the deregulated tyrosine kinase activity of this oncoprotein, but there were no clear differences between wt and dKO cells in commonly activated signal transduction pathways in this type of leukemia cells including Stat5, Erk, p38 or Akt. However, levels of p-Src were increased in the dKO cells.

**Figure 2.**
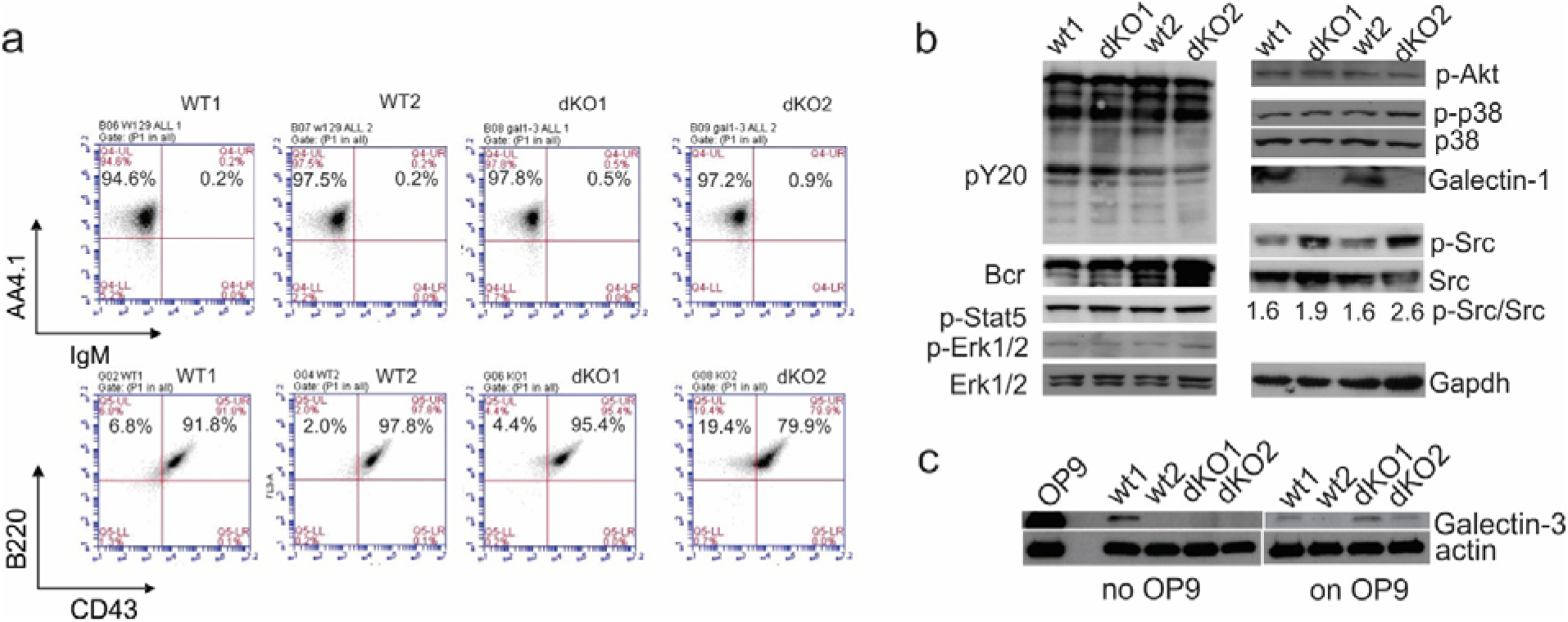
Characterization of murine Bcr/Abl-positive BCP-ALL cells lacking Galectin-1 and Galectin-3 generated by transformation of *Lgals1 x Lgals3 -/-* or matched WT bone marrow cells with p190 Bcr/Abl. Samples include wild type (wt) pre-B ALL #1 and #2, and *Lgals3xLgals1-/-* (dKO) pre-B ALLs dKO1 and dKO2, representing independent transductions. **(a**) FACS analysis for the indicated markers at week 3 after transduction. Gates were set using isotype controls. Numbers: % cells in indicated quadrants. (**b, c**) Western blot analysis with the antibodies indicated to the side of the panels. The ratio of pSrc/Src was determined by densitometric scanning of Western blot film images using Image J software. Gapdh, loading control. BCP-ALL cells grown without stroma. (**c**) Cells as in (**b**) grown alone, or co-cultured for 24 hours with mitomycin-C inactivated murine OP9 bone-marrow stromal cells as a source of Galectin-3.

Endogenous Galectin-3 was only detected in wt1 cells grown alone (Figure 2c) and this was significantly less than the amount of Galectin-3 present in lysates of the murine bone marrow stromal cell line OP9. When the murine BCP-ALL were exposed to Galectin-3 made by OP9 stromal cells through co-culture, three of the four BCP-ALLs endocytosed Galectin-3 detectable by Western blot (Figure 2c).

Physiologically, wt and dKO cells were clearly different. Whereas wild type BCP-ALL cells had robust cell growth, loss of both endogenous Galectin-1 and −3 resulted in considerably reduced proliferation rates (Figure 3a). Using murine BCP-ALL *Lgals3-/-* cells, we previously showed that endogenous expression of Galectin-3 provides protection to the BCP-ALL cells against chemotherapy (21). Using pharmacological inhibition of Galectin-1, we also have evidence that the expression of Galectin-1 provides chemoprotection (23). Therefore, we compared the growth and survival of wt murine BCP-ALL cells to that of the dKO cells under drug treatment.

**Figure 3.**
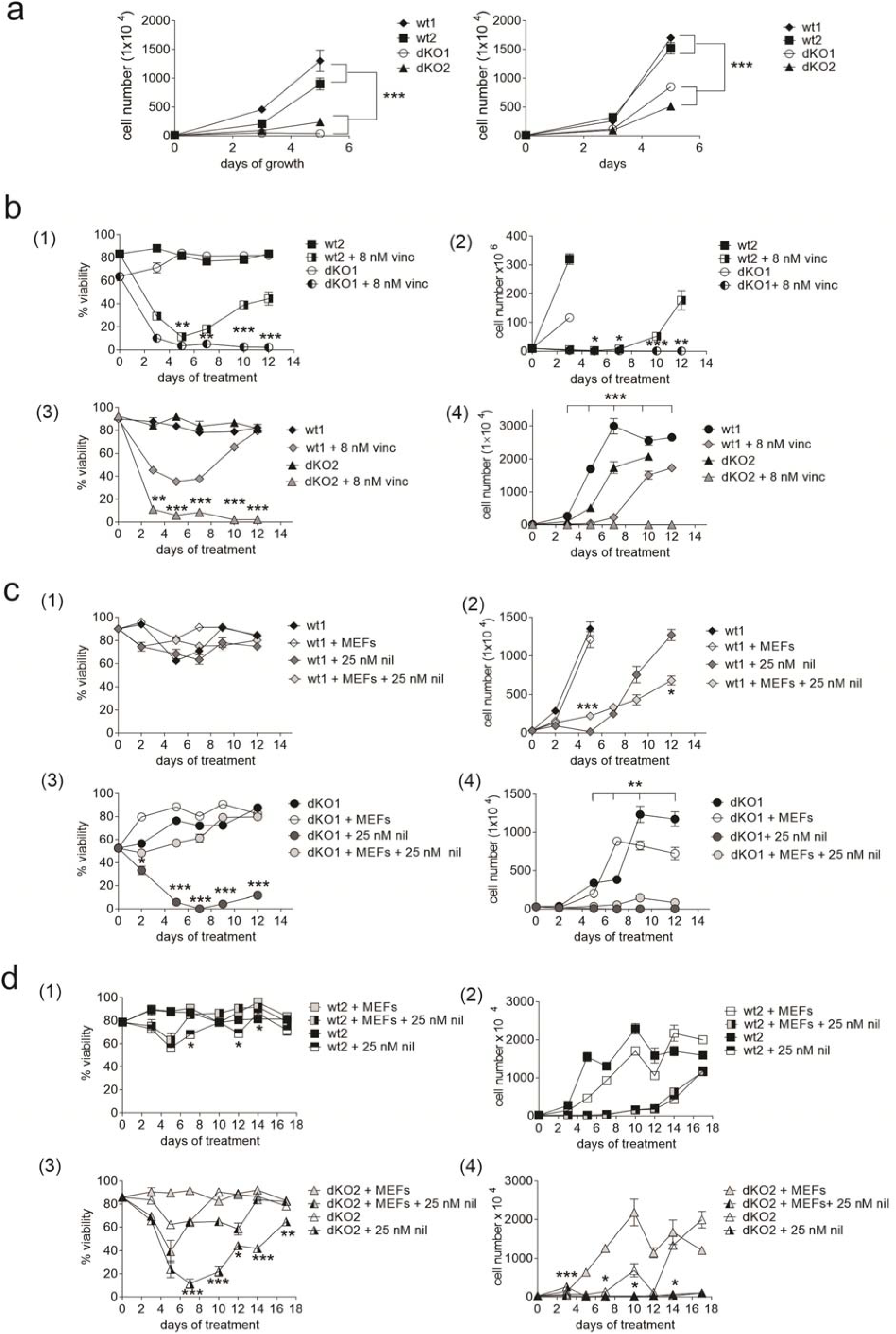
Murine Bcr/Abl-positive BCP-ALL cells lacking Galectin-1 and Galectin-3 have reduced growth and survival. (**a**) Proliferation as measured by viable cell counts of the indicated genotypes. Left panel, 7 weeks after transduction; right panel, 8 weeks after transduction. ***p<0.001 for the indicated group comparisons. (**b-d**) Comparative analysis of viability (left panels) and proliferation (right panels) of dKO and wt ALL cells in a long-term drug treatment (**b**) Cells treated with vincristine. Values for dKO + 8 nM vincristine compared to wt + 8 nM vincristine: *p<0.05, **p<0.01 ***, p<0.001. (**c**, **d**). Nilotinib treatment. Experiments with wt [(1) and (2)] and dKO [(3) and (4)] cells grouped in panels **c** and **d** were performed together but are shown in separate graphs for clarity. Cells at each time point grown without and with MEFs while treated with nilotinib are compared pair-wise at each time point for statistically significant differences. Significant differences are indicated. *p<0.05, **p<0.01 ***, p<0.001

Cells were treated with a non-lethal dose of vincristine, one component of the standard induction chemotherapy regimen for BCP-ALL. As shown in Figure 3b [panels (1) and (3)], wt2 or wt1 BCP-ALL cells treated with vincristine showed an initial drop in viability but resumed proliferation [Figure 3b panels (2) and (4)] as the cells become resistant to the chemotherapy). However, the viability of dKO2 or dKO1 BCP-ALL cells treated with vincristine [Figure 3b panels (1) and (3)] did not recover after 4-5 days of drug exposure and no cell growth was measured [Figure 3b panels (2) and (4)].

Stromal cells (including MEFs, mouse embryonic fibroblasts) provide significant protection to human BCP-ALL cells against many types of drug treatment (14, 15, 17–19). To examine if co-culture with stromal cells could compensate to any degree for lack of endogenous Galectin-1 and Galectin-3 in double null mutant cells, we compared the growth and survival of dKO BCP-ALL cells, cultured with and without MEFs during nilotinib treatment. As shown for wt1 ([Figure 3c, panel (1)], and wt2 [Figure 3d, panel (1)], consistent with their stromal independence, wt BCP-ALL cells had similar viability with or without co-culture with MEFs. Moreover, viability was minimally affected by nilotinib treatment. Nilotinib was cytostatic for the growth of wt1 [Figure 3c, panel (2)]) and wt2 [Figure 3d, panel (2)], and the presence of MEFs had no positive effect on proliferation. For dKO1 [Figure 3c, panel (3)] and dKO2 (Figure 3d, panel (3)], nilotinib treatment decreased their viability, but the presence of MEFs significantly improved this, indicating that some of the cells survived over the duration of the treatment. However, there was still little proliferation [Figure 3c, d panels (4)].

### 2.3. Galectin-1 and Galectin-3 stimulate migration

Stromal cells produce SDF1α, the main chemoattractant for BCP-ALL cells (19, 45). Therefore, when BCP-ALL cells are introduced into a Transwell that has a stromal layer on the bottom, they migrate into the bottom compartment. As shown in Figure 4a, lack of endogenous Galectin-1 and Galectin-3 reduced migration of mouse Bcr/Abl-expressing BCP-ALL cells towards MEFs, indicating that cell-endogenous Galectins contribute to efficient migration.

**Figure 4.**
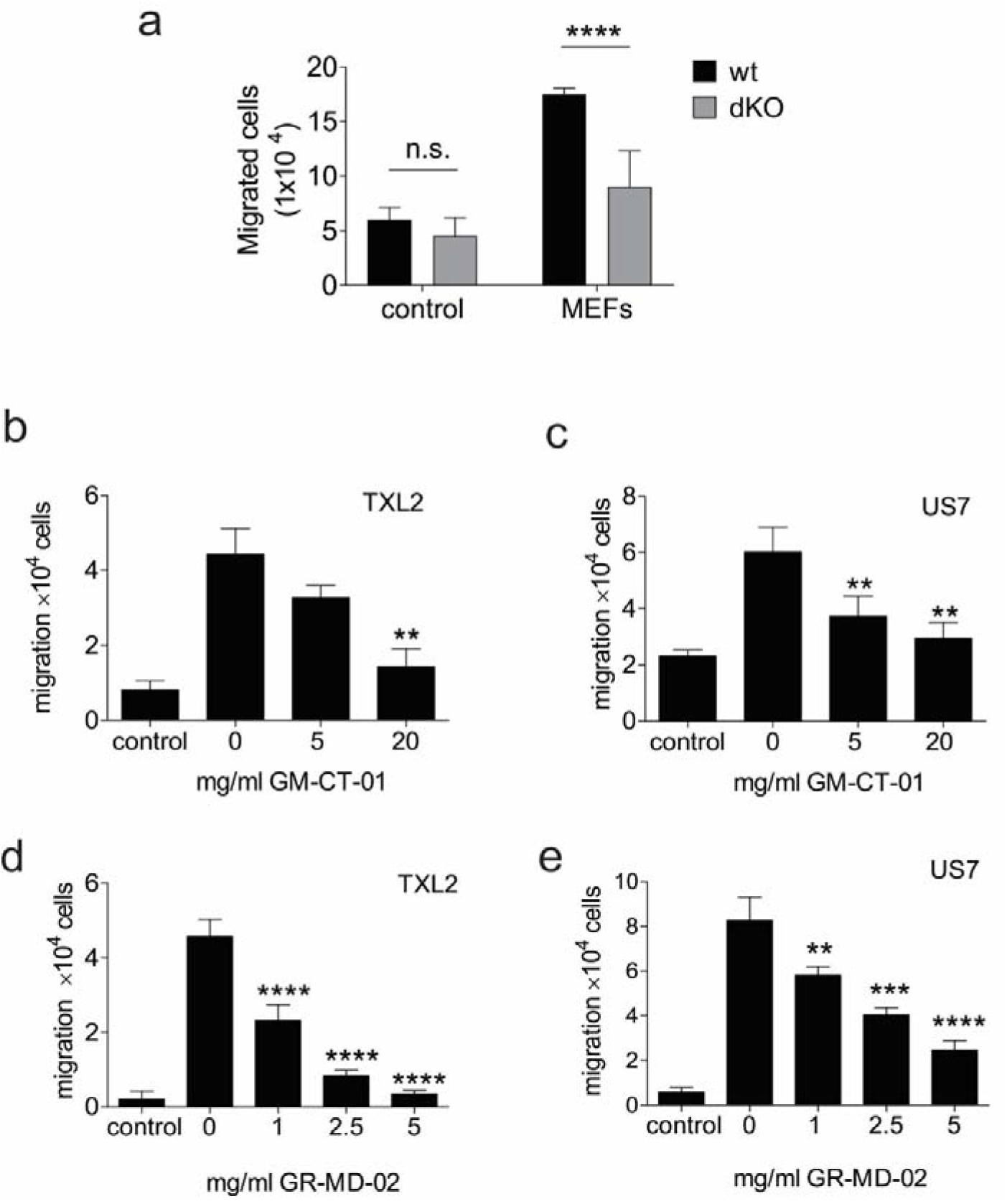
Migration of ALL cells to stroma is promoted by Galectin-1 / Galectin-3. (**a**) mouse BCP-ALL wt1 / wt2 and dKO1 / dKO2 cells were plated in the upper wells of a Transwell and allowed to migrate overnight towards control medium or MEFs plated in the bottom well. ****p<0.001 for differences between combined wt and dKO samples, unpaired t-test. Control samples, not significant. (**b-e**) Overnight migration of human BCP-ALL TXL2 (Ph-positive) or US7 (Ph-negative) to OP9 stromal cells plated in the bottom of Transwells in the absence or presence of the indicated concentrations of GM-CT-01 (**b, c**) or GR-MD-02 (**d, e**). For **b**-**e,** * p<0.05, ** p<0.01, *** p<0.001, ****p<0.0001, one-way ANOVA, between 0 mg/ml compound and samples incubated with the indicated amounts of compound. Controls: only medium, no OP9 cells in bottom well.

To also evaluate the effect of extracellular Galectins on migration, we used a human Bcr/Abl expressing (Ph-positive) ALL, TXL2, in co-culture with OP9 stromal cells as one source of Galectins. We used the galactomannan GM-CT-01 and the galactoarabino-rhamnogalaturonan GR-MD-02 (38, 40) as inhibitors of extracellular Galectin-1/-3. Figure 4b shows that at high concentrations, GM-CT-01 reduced migration of TXL2 towards OP9 stroma. GR-MD-02 (Figure 4d) was active in inhibition of cell motility at lower concentrations than GM-CT-01. The effect of these compounds was not restricted to BCP-ALLs that express Bcr/Abl, as a similar effect was measured in US7, a human BCP-ALL that does not contain the Bcr/Abl oncogene or other known genetic abnormalities (Figure 4c, e).

### 2.4. Intracellular interaction of Galectin-3 with Abl

Actin cytoskeletal reorganization is essential for cell migration, and in some cell types Abl was shown to play a critical role in regulating actin dynamics (46). In leukemias with a *BCR/ABL* translocation, the actin binding domain of Abl is retained in the fusion protein and the Bcr/Abl protein associates with the actin cytoskeleton via the Abl actin-binding domain. Thus Bcr/Abl also regulates cell migration (47, 48). Interestingly, Galectin-3 was shown to form a protein complex with Abl in prostate and breast cancer cell lines and becomes phosphorylated on tyrosine (49–51).

The Ph-translocation is the defining characteristic of chronic myelogenous leukemia (CML), another leukemia that expresses a Bcr/Abl fusion protein. We made use of the CML cell line K562, because it contains multiple copies of the *BCR/ABL* gene (52), to determine if Bcr/Abl and Galectin-3 also interact. Moreover, this cell line also synthesizes Galectin-3 endogenously (24, 53). As shown in Figure 5a (lane total lysate, immunoblotted with Galectin-3 antibodies), we confirmed that K562 cells express Galectin-3 protein endogenously in the absence of stromal cells. Interestingly, Galectin-3 was detected in immunoprecipitates with Bcr/Abl using two different anti-c-Abl monoclonal antibodies (Figure 5a, IP Abl 3F12 or IP Abl Ab-3; bottom panel, Galectin-3 WB). To determine if Galectin-3 becomes tyrosine phosphorylated in these cells, we immunoprecipitated Galectin-3 and used anti-PY20 antibodies to examine the precipitated protein. Figure 5b (right panel: IP Gal3, IB pY20) shows that Galectin-3 is tyrosine phosphorylated and confirmed it is detected in the Bcr/Abl immunoprecipitate (left panel: IP 3F12, IB: Gal3)).

**Figure 5.**
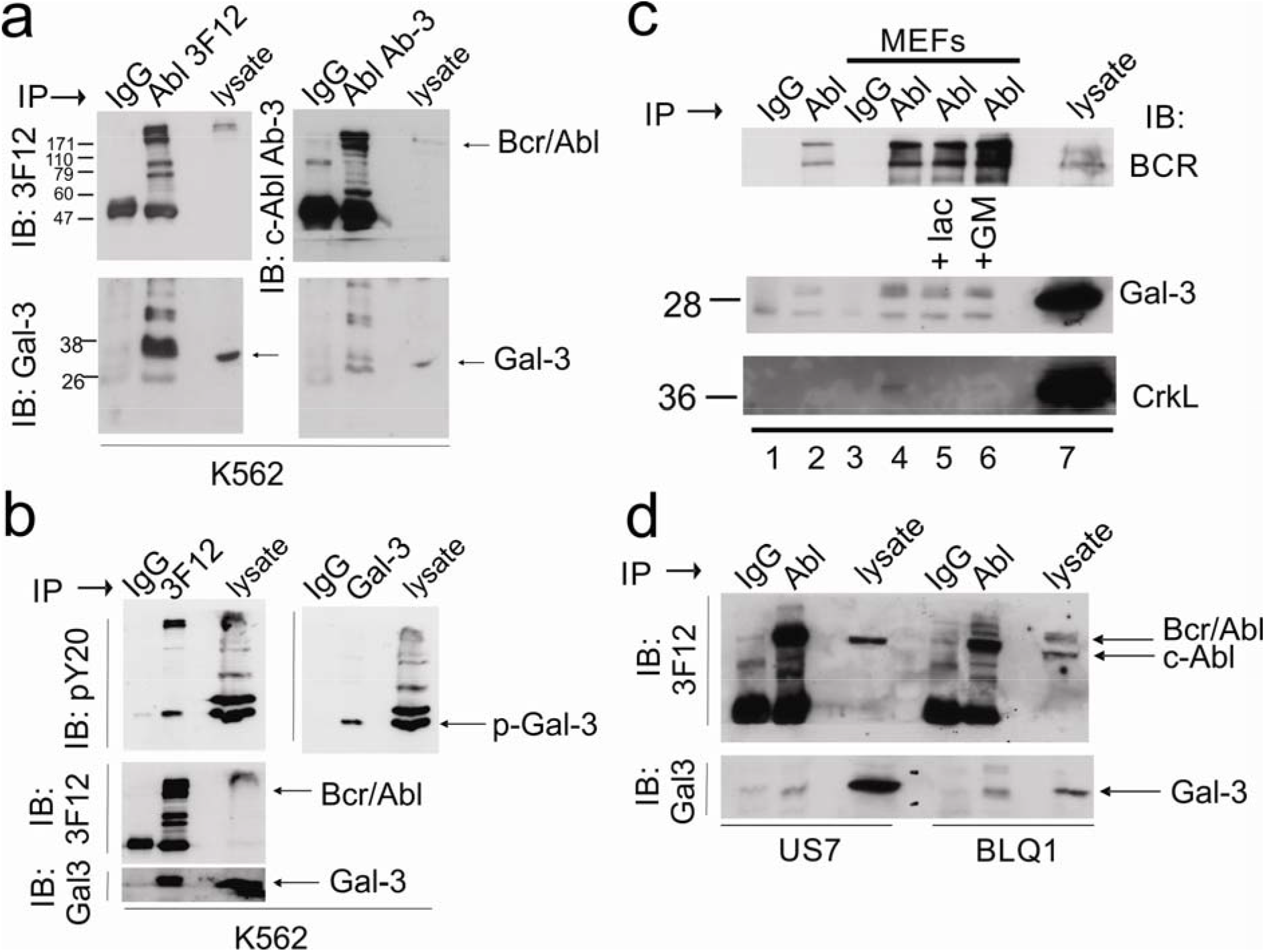
Complex formation of Galectin-3 with (Bcr/)Abl in leukemia cells. Lysates made from (**a-c**) the CML cell line K562 or (**d**) BCP-ALLs. **(a).** Immunoprecipitation with control Ig or two different anti-c-Abl antibodies: 3F12 or c-Abl Ab-3 (Calbiochem) as indicated above the lanes. Antibodies used for immunoblotting are indicated to the left. Arrows to the right of the panels point to the location of Bcr/Abl and Galectin-3 (Gal-3). Note: the right upper panel was exposed longer than the left upper panel. (**b**). Immunoprecipitation with anti-Abl 3F12 or Galectin-3 antibodies followed by immunoblotting with the antibodies indicated to the left of the panels. (**c**) Immunoprecipitation from K562 lysates of cells grown as suspension culture (lanes 1-2) or on MEFs (lanes 3-6) as indicated above the panel. Lane 7, total cell lysate. Antibodies used for IP include control IgG (lanes 1 and 3) or 3F12 anti-Abl (lane 2, lanes 4-6). Lane 5, cells treated for 2 hours with 50 mM lactose; lane 6, cells treated with 10 mg/ml GM-CT-01 for 2 hours. After immunoblotting with Galectin-3, the panel was stripped and re-probed with anti-Crkl antibodies. A 12% SDS-PAA gel was used to obtain separation in the lower molecular weight range. (**d**) Immunoprecipitation of c-Abl and Bcr/Abl from lysates of human BCP-ALL US7 (Ph-negative) and BLQ1 (Ph-positive). Cells were grown on OP9 stroma before preparation of lysates.

Because Galectin-1 and Galectin-3 are well-conserved, most antibodies recognize both human and murine proteins. However, human and mouse Galectin-3 differ in molecular weight [Figure S1b and (24)], and thus we are able to distinguish the two sources in some co-cultures. A Western blot for Galectin-3 in total lysates of BCP-ALL cells is expected to provide information on the intracellular Galectin-3 and that which is strongly cell-surface bound. We found that GM-CT-01 clearly reduced the amount of the stromal-produced Galectin-3 (top Galectin-3 band) associated with the leukemia cells (also see Figure S1b, right panel, 24 hrs), but it had less effect on the Galectin-3 that had been made by the leukemia cells (lower Galectin-3 band). This is consistent with the concept that such compounds, because of their relatively large size, mainly have an effect on displacement of extracellular Galectins. Also, GM-CT-01 treatment did not seem to affect Galectin-1 levels in K562 lysates (Figure S1b left panel.

To investigate if exogenous Galectin-3 provided by stroma also interacts with Abl, we co-cultured the K562 with MEFs, prepared lysates, and preformed immunoprecipitations with anti-Abl 3F12 antibodies. As shown in Figure 5c, compared to K562 cultured alone, the co-cultured cells contained a higher molecular Galectin-3 weight band (compare lanes 2 and 4, Gal-3 WB) representing murine Galectin-3. Mouse Galectin-3 as well as endogenous human Galectin-3 co-immunoprecipitated with Bcr/Abl (Figure 5c, double band in IPs from K562+MEF lysates, 3F12 IPs, Galectin-3 WB). Re-probing of the membrane with anti-Crkl antibodies showed that Crkl had also been co-immunoprecipitated with Bcr/Abl as expected (54). These results suggest that both endogenously produced as well as exogenous Galectin-3 bound to or taken up by K562, can interact intracellularly with (Bcr)/Abl.

To confirm that Bcr/Abl indeed forms a complex with exogenously produced Galectin-3, we immunoprecipitated Bcr/Abl from human Ph-positive BLQ1 ALL cells. These do not express much Galectin-3 endogenously under steady state conditions (22). Figure 5d (IP 3F12 IB: Galectin-3) shows that we also detected co-immunoprecipitation of Bcr/Abl with Galectin-3 in BLQ1 ALL cells, indicating that mouse Galectin-3 and human Bcr/Abl also interact. As we also detected co-immunoprecipitation of c-Abl with Galectin-3 in US7 cells, which are Ph-chromosome negative and do not contain Bcr/Abl (Figure 5d), interaction of Abl with Galectin-3 may be a common mechanism that could regulate migration of BCP-ALL cells.

### 2.5. Effect of GM-CT-01 or GR-MD-02 on ALL signaling

OP9 stromal cells synthesize and secrete high levels of Galectin-3 (22). As mentioned, our studies have shown that under non-stressed conditions, the Galectin-3 on and inside BCP-ALL cells mainly originates from the OP9 stromal cells in the tissue co-culture model (22, 24). We therefore investigated if GM-CT-01 could reduce binding of Galectin-3 to BCP-ALL cells. We harvested ALL cells from underneath the stroma and stained the ALL cells for Galectin-3 with or without prior incubation with GM-CT-01. As shown in Figure 6a, binding of Galectin-3 antibody to the cells was reduced in the presence of GM-CT-01, providing evidence that the compound is able to decrease cell surface-bound Galectin-3 on these cells.

**Figure 6.**
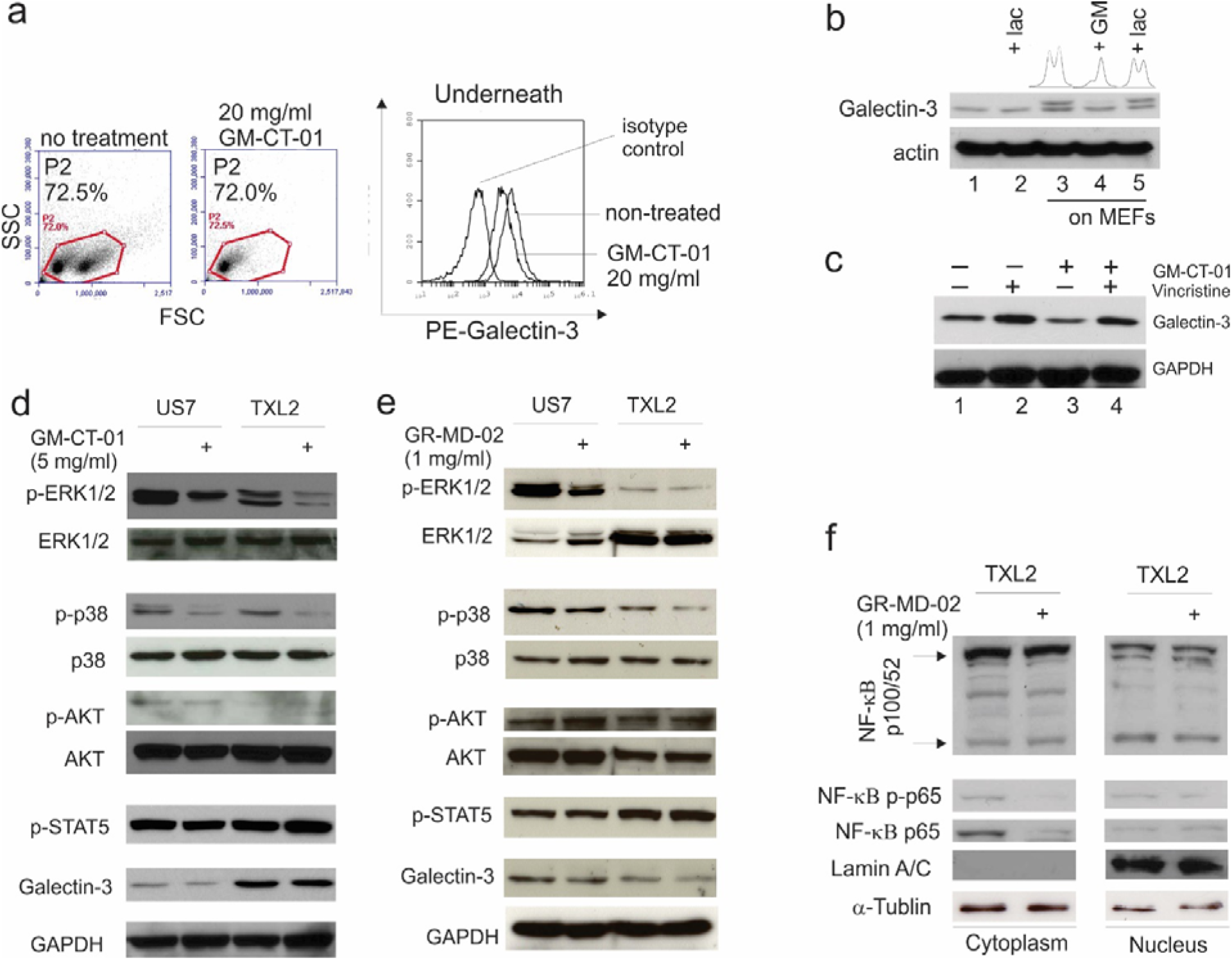
GM-CT-01 and GR-MD-02 attenuate mitogenic signaling in ALL cells. (**a**) ALL cells in co-culture with OP9 cells were treated with 20 mg/ml GM-CT-01 for 2 hours. ALL cells from underneath the stroma were assayed for cell surface expression of Galectin-3 using FACS: left, gating; right Galectin-3 cell surface positive cells in gate P2. Control, ALL cells underneath OP9 not treated with GM-CT-01. (**b**) Western blot analysis for Galectin-3 expression in K562 cells grown in suspension or plated on MEFs. Lanes 2 and 5, cells treated for 2 hours with 50 mM lactose (lac); lane 4, cells treated for 2 hrs with 20 mg/ml GM-CT-01 (GM). The histograms above lanes 3-5 represent densitometry using Image J. (**c**) Western blot analysis for Galectin-3 in US7 cells on day 20 of treatment with no drugs, with 2.5 nM vincristine, with 1 mg/ml GM-CT-01 or a combination, as indicated above the panel, while co-cultured with OP9 stroma. (**d**) Western blot analysis of US7 cells and TXL2 total cell lysates after incubation with 5 mg/ml GM-CT-01 for 72 hours in the presence of OP9 stromal cells. Blots were stripped and re-probed with Erk1/2, p38, Akt and Gapdh as loading controls. (**e, f**) US7 cells and TXL2 cells were incubated with 1 mg/ml GR-MD-02 for 72 hrs in the presence of OP9 cells. Blots were stripped and reprobed with Erk1/2, p38, Akt and Gapdh as loading controls. (**f**) NF-κB p100/52, RelA NF-κB p-p65 and NF-κB p65 WB on TXL2 cytoplasmic and nuclear fractions. Lamin A/C and α-tubulin, loading controls for nuclear and cytoplasmic fractions, respectively.

To compare the effect of GM-CT-01 on extracellular and intracellular Galectin-3, we generated lysates of K562 alone or in co-culture with MEFs, after treatment, or not, with GM-CT-01. As shown in Figure 6b, when these cells are plated on MEFs and the samples are run on a lower percentage SDS-PAA gel, an additional band is visible with a slightly larger molecular mass, representing murine Galectin-3. Interestingly, treatment with GM-CT-01 reduced the signal from the murine Galectin-3 (Figure 6b, lane 5; also see Figure 5c lane 6 panel Gal3 immunoblot), supporting the proposed reduction by this compound of extracellular Galectin-3 bound to leukemia cells. 50 mM lactose as a competitor had minimal effects on the endogenous Galectin-3 in K562 (Figure 5c lane 2). Compared to 20 mg/ml GM-CT-01, it also did not reduce mouse Galectin-3 detected in the K562 lysates (Figure 5c lane 5 and Figure 6b lane 5).

We have previously shown that BCP-ALL cells treated with chemotherapy including vincristine also produce Galectin-3 endogenously (22), as illustrated in Figure 6c (compare lane 1, no treatment to lane 2, vincristine treatment). Interestingly, GM-CT-01 treatment reduced Galectin-3 levels both in non-vincristine as well as vincristine-treated cells (lanes 3 and 4). We next analyzed the possible consequences of this treatment on intracellular signaling. Figure 6d shows that a 3-day treatment with GM-CT-01 attenuated pErk1/2 signals in both US7 and TXL2 cells and also reduced endogenous levels of p-p38. We also treated cells with GR-MD-02. As shown in Figure 6e, similar to GM-CT-01, the phosphorylation of p38 and Erk1/2 was reduced by this compound, whereas Akt and STAT5 phosphorylation were not visibly affected. Cytoplasmic NF-kB p-p65 and NF-kB p65 levels were also decreased after long-term exposure to GR-MD-02 (Figure 6f).

### 2.6. GM-CT-01 or GR-MD-02 inhibit proliferation of ALL cells in the presence of stromal support

We examined whether GM-CT-01 alone or in combination with therapeutic drugs affects the viability of ALL cells in the presence of OP9 stromal cells. As shown in Figure 7a, whereas GM-CT-01 alone had no obvious effect on TXL2 ALL cells, the addition of GM-CT-01 to nilotinib-treated cells further decreased cell viability and cell numbers beyond nilotinib mono-treatment. The compound also significantly reduced US7 ALL cell viability beyond that of vincristine mono-treated cells (Figure 7b) and had a visible effect on US7 leukemia cells when combined with vincristine (Figure 7c).

**Figure 7.**
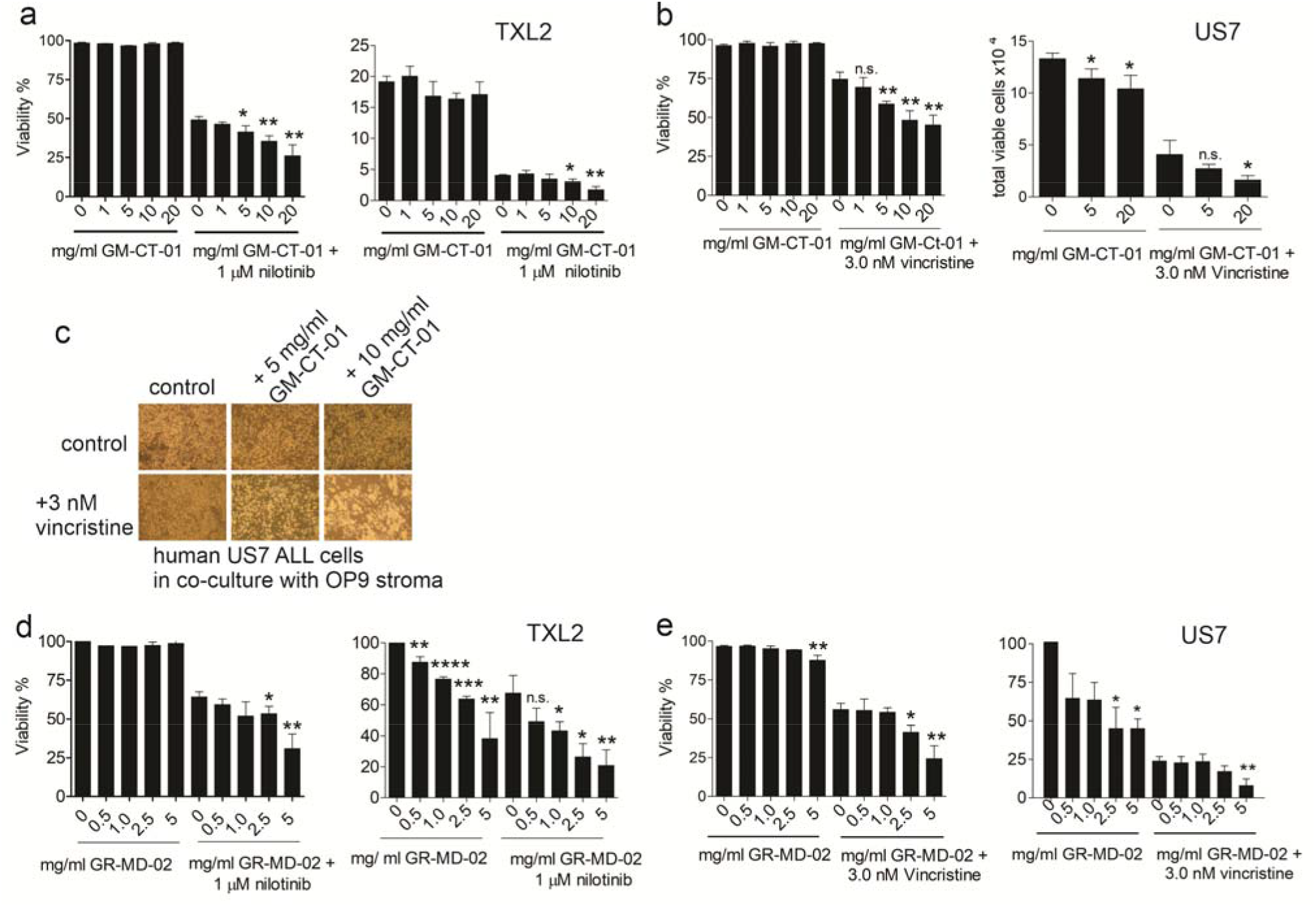
Combination drug treatment with GM-CT-01 or with GR-MD-02 inhibits proliferation of ALL cells. (**a**) Viability (left panel) and cell counts (right panel) of Ph-positive TXL-2 ALL cells treated with GM-CT-01 and nilotinib for 96 hours in the presence of OP9 cells. (**b**) Viability (left panel) and cell counts (right panel) of US7 cells treated for 96 hours with GM-CT-01 alone or in combination with vincristine in the presence of OP9 cells. (**c**) Phase contrast images of US7 cells co-cultured with irradiated OP9 stroma and treated with GM-CT-01 and vincristine as indicated. (**d, e**) Viability and cell counts of TXL2 cells (**d**) or US7 (**e**) treated with GR-MD-02 or a combination treatment as indicated. Viable cells and cell numbers determined using Trypan blue exclusion. * p<0.05, ** p<0.01, ***p<0.01, ****p<0.0001 compared to 0 mg/ml samples, representing treatment with nothing or only with nilotinib or vincristine, unpaired t-test. Control samples, DMSO at the same final concentration as the vincristine/nilotinib samples.

We additionally evaluated GR-MD-02. This compound was effective at lower concentrations than GM-CT-01. Similar to GM-CT-01, the compound had little effect on the viability of the cells as monotreatment (Figure 7d, e, left panels GR-MD-02 alone) but did inhibit proliferation of both TXL2 (Figure 7d, right panel) and US7 (Figure 7e right panel) as measured by a reduction in the total viable cell count. Nilotinib as tyrosine kinase inhibitor is not extremely cytotoxic, but its combination with relatively lower amounts of GR-MD-02 (Figure 7d, right panel, 1 mg/ml and lower) produced a significant reduction in cell counts. GR-MD-02 with vincristine (Figure 7e left panel) also had increased toxicity at 2.5 and 5 mg/ml compound and a cytostatic effect on the US7 ALL cells, at 5 mg/ml (Figure 7e, right panel).

## 3. Discussion

### 3.1 Overlapping mRNA expression of Galectin-3 and Galectin-1

Galectin-1 and Galectin-3 have each been widely studied in the context of cancer and as immune cell modulators but relatively little is known about their functional redundancy when expressed in the same cells. Concurrent expression is relatively common: for example, normal murine B-cell progenitors express both *Lgals1* and *Lgals3* (Figure S2) although clearly the mRNA expression of *Lgals1* is higher than that of *Lgals3* in the Hardy fraction precursors (Figure S2a). *Lgals1* expression appears to be developmentally regulated, with decreasing levels as the cells progress with normal maturation, whereas *Lgals3* does not show such correlation. Interestingly however, quiescent murine hematopoietic stem cells do express high levels of endogenous Galectin-3 and this is functionally important to downregulate cell cycle (55). Normal human bone marrow stromal cells (Figure S3a) and OP9 cells express both Galectins, but only *Lgals3* is affected by imatinib treatment (Figure S3b).

When pediatric ALL patients are treated with chemotherapy over the course of 33 days, expression of both *LGALS1* and *LGALS3* mRNA significantly increases in total bone marrow [Figure S4a; *LGALS3* graph also shown in (24)]. Ebinger *et al* (56) compared gene expression of dormant non-proliferating BCP-ALL cells with that of actively dividing cells in the same population and derived a 250-gene signature characteristic of the former. Galectin-1 but not Galectin-3 is included in this gene set. The same authors also compared diagnosis to MRD samples after 33 days of chemotherapy, and in that data set Galectin-3 but not Galectin-1 was significantly upregulated in the MRD samples. Similarly, other hematopoietic cell types show concurrent but differentially regulated Galectin-1 and Galectin-3 expression. Compared to normal CD34+ progenitor cells, CD34+ AML cells have higher *LGALS1* levels, whereas CD34-AML cells express elevated *LGALS1* and *LGALS3* (Figure S4b). NK cells from healthy donors and from patients with multiple myeloma have significantly higher *LGALS1* but not *LGALS3* expression when they are expanded and activated by *ex vivo* co-culture (Figure S4c).

### 3.2 Overlapping function of Galectin-1 and Galectin-3

Overlapping expression patterns of Galectin-1 and Galectin-3 raise the question of functional redundancy. Indeed, Galectin-1 and Galectin-3 have many binding partners in common, such as von Willebrand factor, the lysosomal protein Lamp1 (57–59) and the mature B-cell transcription factor OCA-B (60), suggesting that they may have overlap in function. In human mesenchymal retinal pigment epithelial cells, many other glycoproteins were identified that bind both Galectin-1 and Galectin-3 (61). In agreement with functional overlap, simultaneous knockout of Galectin-1 and Galectin-3 in MEFs strongly reduced EGF-stimulated activation of K-Ras, Erk and Akt, compared to Galectin-1 or Galectin-3 single null mutant MEFs (62). In our studies using a different cell type we did not detect differences in baseline levels of activated Erk or Akt in *Lgals1 x Lgals3 -/-* dKO BCP-ALL cells compared to wt. This could be explained because we studied cancer cells, which express Bcr/Abl as driver oncogene, obviating the need for exogenous cytokine signaling and making the cells IL-7 independent, while activating multiple endogenous signal transduction pathways (8). Interestingly, we did detect increased phosphorylation of an activating tyrosine residue common to a number of Src-family kinases (SFK) such as Lck and Lyn, in the double knockout cells. These antibodies are directed against Y419 in Src but may also react with Y394 in Lck, or Y397 in Lyn, both of which are also highly expressed in BCP-ALL. This residue can be dephosphorylated by the transmembrane phosphatase CD45 (63, 64). Although it will require further investigation, the specific loss of Galectin-1 could be the cause of increased SFK phosphorylation since Galectin-1 is known to regulate the basal and activating signaling of such SFK through binding to the extracellular domain of CD45 and regulating its phosphatase activity towards SFK as reported in T cells (65, 66).

Few studies have examined the functional consequences of simultaneous genetic ablation of Galectin-1 and Galectin-3. The original report documented that concurrent loss of *Lgals1* and *Lgals3* function by gene targeting in mice is compatible with relatively normal development (67), indicating that these genes do not contribute critically to steady-state physiology. Using double knockout cells, Clark *et al* showed that Galectin-1 and Galectin-3 both regulate mature B-cell functions in a mouse model of autoimmune disease (68). Also, Sirko *et al* showed that Galectin-1 and Galectin-3 together promote astrocyte reactivity, and that exogenously added Galectin-3 but not Galectin-1 can normalize the double mutant phenotype. In our study we found that the double knockout Galectin-1 and Galectin-3 murine BCP-ALL cells proliferated significantly less well than wild type controls, indicating that these cells clearly are defective in some aspects of mitogenic signaling, or have other endogenous deficiencies related to, for example, cell cycle progression. It is possible that the proliferation defect of the dKO BCP-ALL cells also causes the greater vulnerability to nilotinib or vincristine monotherapy. Interestingly, the proliferation defects of the dKO BCP-ALL cells were mitigated by co-culture with a fibroblast stromal layer, and their ability to withstand drug treatment was also enhanced by the presence of these cells. Therefore, it is likely that the stromal layer is providing some of the missing functions, which could include the production and secretion of Galectin-3 and Galectin-1 (22).

### 3.3. High molecular weight compounds to inhibit all extracellular Galectin-1 and Galectin-3

Our co-culture system of leukemia cells with stromal cells is intended to model the bone marrow microenvironment, where leukemia cells are also protected and supported. In this relatively more complex system, there are two sources of extracellular Galectin-1 and Galectin-3: (1) that which is secreted by stromal cells, can bind to the surface of the leukemia cells and become endocytosed, and (2) that which is produced endogenously by the leukemia cells, of which an unknown fraction is transported to the cell surface and/or secreted. If any of these sources of extracellular Galectin plays an especially critical role in this system is difficult to tease out: Galectin-3 levels may be dynamically regulated by the presence of cellular stress such as chemotherapy whereas for Galectin-1, the developmental stage of the precursor B-cell may be the most important factor that determines expression levels. In fact, Galectin-1 levels are higher in more primitive B-cell precursors, and in addition a specific stromal cell type has been identified in the bone marrow that produces Gaelctin-1 and regulates early B-cell development (69), as further reviewed in (70).

It is generally assumed, that an important extracellular function of Galectin-3 is to bind poly-LacNAc modified glycoproteins, which is needed to promote lattice formation and stimulate subsequent receptor clustering. This is known to enhance intracellular signal transduction. In concordance with this, in our study, blocking of Galectin-3/Galectin-1 by GM-CT-01 or GR-MD-02 down-regulated levels of p-Erk1/2 and p-p38, but no effects on the Akt pathway were seen. In contrast, in myeloma cells, treatment with a different compound, the pectin-derived GCS-100 reduced activation of Akt (71). Thus, the effects of compounds such as GM-CT-01 or GR-MD-02 may depend on which target glycoproteins are bound by Galectin-3 or Galectin-1 on different cell types.

We found that both GR-MD-02 and GM-CT-01 inhibited activation of the Erk pathway in Bcr/Abl-positive and non-Bcr/Abl-transformed human ALL cells, suggesting that such compounds have a general effect such as interfering with the migration of the BCP-ALL cells to the stroma and/or reducing contact with it. Chauhan *et al* (72) observed that GCS-100 overcame bortezomib resistance and enhanced dexamethasone-induced apoptosis in multiple myeloma cells. Streetly *et al* (71) reported similar results. This is consistent with our data which show that such compounds in combination with a therapeutic drug additionally enhance the effects of those drugs in the presence of microenvironmental protection *in vitro*.

We used GR-MD-02 and GM-CT-01 as dual Galectin-1 / Galectin-3 inhibitors based on *in vitro* data reporting a similar K_d_ for GM-CT-01 and GR-GM on Galectin-1 and Galectin-3 (35). However, both compounds have also been used as if they mainly inhibit extracellular Galectin-3 [GM-CT-01 (73); GR-MD-02 (74–77)]. Whether this is a reasonable assumption is not clear: GM-CT-01 interacts with a Galectin-1 domain not including the classical carbohydrate-binding site, but this is *in vitro* (38). Moreover, Stegmayr *et al* (78) reported an IC50 of 4 mg/ml for Galectin-3 and >20 mg/ml for Galectin-1 using GM-CT-01 in an *in vitro* inhibitory binding assay to asialofetuin. These high concentrations are in agreement with the concentrations of GM-CT-01 needed to obtain a biological effect in our studies and raise the concern for off-target effects.

Thus, although the use of these compounds supports the premise that inhibition of the interaction of stromal-produced Galectin-3 and Galectin-1 is useful to chemosensitize BCP-ALL cells, a need remains for the development of more specific and potent inhibitors which include anti-Galectin-3 antibodies (79), novel carbohydrate mimetics [Bum-Erdene *et al;* in preparation and (24)] or peptides [Battacharya *et al*, in preparation]. In addition, because drug-induced Galectin-3 protects BCP-ALL cells, inhibitors that are able to enter cells and interfere with Galectin-3 activity at that location would be needed.

### 3.4 Conclusion

Overall, our studies suggest that the homing of ALL cells to the close proximity of protective stromal cells, such as are present in the bone marrow, is important and may result in the persistence of drug-insensitive cells that can give rise to relapse. Thus carbohydrate-based drugs that interfere with the binding of stromal-produced Galectin-3 and Galectin-1 to the surface of BCP-ALL cells may prevent firm adhesion contacts from forming between the two cell types and sensitize the ALL cells to cytotoxic drugs. Our previous results showed that both endogenous and stromal-produced Galectin-3 protect human BCP-ALL cells against chemotherapy (21, 24), and that pharmacological inhibition of Galectin-1 using PTX008 also sensitizes such cells to drugs (23). Therefore, a strategy of combined inhibition of both Galectin-1 and Galectin-3 function, both extracellularly and intracellularly, using small molecule inhibitors, would be optimal to chemo-sensitize BCP-ALL cells.

## 4. Materials and Methods

### 4.1. Gene expression analysis

Meta-analysis of *LGALS3* and *LGALS1* (Galectin-3 and Galectin-1) expression of RNAs from pediatric Ph-positive ALL at diagnosis was performed on GEO Datasets accession GSE28497 and GSE79533, described in (41) and in (42), respectively. Processed data in the series matrix files represent values normalized by MAS5.0 and baseline transformed to a median target intensity. Txt file values for *LGALS3* and *LGALS1* (probe sets 208949_s_at and 201105_at) imported into Excel were manually extracted into Prism5.0.

Mice transgenic for the human P190 form of Bcr/Abl (Jackson Labs strain 017833) develop precursor B-lineage (pre-B) acute lymphoblastic leukemia, on average within 3 months of birth, when on a C57Bl/6J background. Bone marrows were isolated from control C57BLl/6J mice and from transgenic Bcr/Abl mice when they had not yet developed full-blown leukemia (<60 days of age), from Bcr/Abl transgenic mice with overt leukemia and packed bone marrows (>90 days of age), and from fully leukemic mice that had received a seven-day treatment with 75 mg/kg AMN107 (nilotinib). Three mice were used per condition, and cells from each mouse were processed separately. Pre-B cells from these twelve bone marrows were flow-sorted using CD19 and AA4.1 as markers. Total RNA from CD19+ AA4.1+cells used for microarray analysis was isolated by RNeasy (QIAGEN) purification. Double-strand complementary DNA was generated from 5 μg of total RNA using a poly(dT) oligonucleotide that contains a T7 RNA polymerase initiation site and the SuperScript III reverse transcription (Invitrogen). Biotinylated cRNA was generated and fragmented according to the Affymetrix protocol and hybridized to 430 mouse microarrays (Affymetrix). As described in Trageser *et al* (44), Cel files from GeneChip arrays were imported to the BRB Array Tool (http://linus.nci.nih.gov/BRB-ArrayTools.html) and processed using the RMA algorithm (Robust Multi-Array Average) for normalization and summarization. Data are available at https://www.ncbi.nlm.nih.gov/geo/query/acc.cgi?acc=GSE110104

### 4.2. Drugs and reagents

GM-CT-01 and GR-MD-02 were provided by Galectin Therapeutics, Inc (Norcross, Georgia, USA) and were stored at 4°C. Nilotinib (AMN107) was obtained from Novartis (Basel, Switzerland). Nilotinib was dissolved in DMSO and stored at −20°C. A vincristine sulfate solution was obtained from Hospira Worldwide Inc. (Lake Forest, IL, USA).

### 4.3. Treatment with drugs, cell proliferation and viability, flow cytometry

For proliferation assays, US7 or TXL2 cells were cultured in a 96-well plate at a density of 5×10^4^/well, in the presence of irradiated OP9 cells. Cells were treated with GM-CT-01 or with GR-MD-02, in combination with nilotinib or vincristine. Controls for nilotinib or vincristine were DMSO at the dilution matching the drug samples. After drug exposure, cells were collected and re-suspended in culture medium containing 0.1% (wt/vol) Trypan blue. Trypan blue-excluding and total cells were counted using a hemocytometer. All drug sensitivity assays were done in triplicate wells. Viability of the cells is expressed as the percentage of Trypan blue-excluding cells of the total number of cells. Data points show the mean ±SEM of triplicate samples.

To assay for the ability of GM-CT-01 to displace Galectin-3 using FACS, TXL2 cells were harvested from underneath the OP9 stromal layer. Cells were incubated with or without 20 mg/ml GM-CT-01 for 2 hrs at 37°C in complete medium, then washed with PBS. DTSSP was added at 5 mM to cross-link extracellular Galectin-3 to the cell surface. After a 30-min incubation at RT, the reaction was terminated by a 15-min incubation with 50 mM glycine at pH 7.5. After a wash in PBS, cells were incubated with PE-Galectin-3 (Biolegend cat# 126706) or matched isotype control for 15 mins and analyzed on BD Accuri C6 cytometer (BD Biosciences, San Jose, CA).

FACS analysis of mouse wt and dKO BP ALL cells used antibodies against AA4.1 (eBiosciences cat#17-5892-82), CD43 (BD cat#553270) and CD45/B220 (Biolegend cat#103132). Gates were set based on isotype controls.

### 4.4. Migration Assay

For migration assays, human TXL2 or US7 cells (1×10^5^) were seeded into the upper well of Transwell plates with a 5 μm pore size. The lower chamber contained a layer of irradiated OP9 stromal cells with different concentrations of GR-MD-02 or GM-CT-01 as indicated. Wells without stromal cells in the bottom chamber served as controls. ALL cells migrated to the bottom wells were counted after overnight incubation. Migration of wt1, wt2, dKO1 and dKO2 BCP-ALL cells was measured individually, but values of wt1/wt2 and dKO1/dKO2 were combined for statistical analysis in Figure 4a.

### 4.5. Cells

Murine BCP leukemia: Murine leukemia cells were generated from bone marrows of age and sex matched wild-type (129 P3/J *Lgals1 x Lgals3 ^+/+^*) controls and Galectin-1/Galectin-3 (129 P3/J *Lgals1 x Lgals3 ^-/-^*) double knockout mice on a C57Bl/6J background. Bone marrow cells were transduced with P190 Bcr/Abl-encoding retroviruses as previously described (18). In brief, transduced bone marrows were grown for 5 days with 10 ng/ml rmIL-7 and cultured in IMDM with 50 μM β-mercaptoethanol in 20% FBS, 1% L-glutamine, 1% penicillin/streptomycin. The murine BCP-ALL cells were grown without stroma except where indicated. Stroma, when used, includes mouse embryonic fibroblasts (MEFs) or mitotically inactivated (irradiation or mitomycin treated) OP9 cells. All assays were performed within ≈ 6 months of the initial transduction.

Cell lines and PDX-derived human BCP-ALLs: The murine OP9 stromal cell line (CRL-2749) and the human CML cell line K562 were obtained from the ATCC (Manassas, VA, USA). Human ALL cells including Ph-positive TXL2 and BLQ1, and Ph-negative US7 cells were described previously (17). Human leukemia cells were grown in MEM-α medium supplemented with 20% FBS, 1% L-glutamine and 1% penicillin/streptomycin (Invitrogen Corporation). Human ALL cells were cultured in the wells of a 6-well plate or a 96-well plate at a density of 0.5-1×10^6^ cells/ml, in the presence of irradiated OP9 cells.

Mouse embryonic fibroblasts (MEFs): E13.5 embryos were isolated from timed matings of C57Bl/6J mice. Internal organs and heads were removed and tissue homogenized by mincing with razor blades and pressure from the back of a syringe (80–82). Single cell suspensions were further generated by a 30-min incubation at 37 °C in 5 ml Trypsin-EDTA. Tissue chunks were removed with a cell strainer, and cells were plated at a density of 6×10^6^ per 15 cm dish in DMEM + 10% FBS, P/S, and L-glutamine.

### 4.6. Western blotting and immunoprecipitation

Anti c-Abl Ab-3 antibodies for immunoprecipitation were from Calbiochem (San Diego CA). For immunoprecipitations, 1 mg of protein was used. Lysates were pre-cleared by incubation with 3 μg rabbit or mouse IgG and PAA. ALL cells were lysed for 30 minutes on ice in RIPA buffer (50 mM Tris-HCl, pH 8.0, 150 mM NaCl, 1% Triton X-100, 0.5% deoxycholate, 0.1% SDS, 5 mM EDTA) containing PMSF, aprotinin, leupeptin, pepstatin A, Na-fluoride and Na-orthovanadate. For detection of NF-kB (p100/52, p65), a nuclear extraction kit (Imgenex) was used to separate nuclear and cytoplasmic fractions. Cell extracts were subjected to 8-15% sodium dodecyl sulfate-polyacrylamide gel electrophoresis. Antibodies used include pY20 (BD-Transduction San Jose CA), Galectin-3 (Biolegend, San Diego, USA), phospho-ERK1/2, phospho-p38, phospho-STAT5, phospho-AKT, AKT, (Cell Signaling Technology, USA), ERK1/2, NF-kB p65, Bcr N-20 (Santa Cruz Biotechnology, USA), NF-kB p100/52 (Millipore, USA) using standard procedures. Gapdh (Chemicon International, USA) or β-actin (Sigma, USA) antibodies were used as a loading control. The phospho-SFK (Y416, Cell Signaling Technology) antibody recognizes pY416 in Src but may cross-react with other Src family members (Lyn, Fyn, Lck, Yes and Hck) when phosphorylated at equivalent sites.

### 4.7 Statistical analysis

Statistical tests are mentioned in the figure legends and were done using GraphPad Prism versions 5-9. A False Discovery Rate approach was adopted to decide the significance of the comparison based on that no more than 5% of the discoveries will be false discoveries. Pearson correlation analysis was performed to examine expression correlations between *LGALS1* and *LGALS3* genes in public data sets. A two-tailed p-value was calculated to suggest the significance of the correlation and a p-value < 0.05 was used to decide whether the two sets of expression values were correlated.

## Supporting information

previous peer review

## Acknowledgments

We very gratefully acknowledge Francoise Poirier for the wt and *Lgals1 x Lgals3 -/-* bone marrow samples, without which this study would not have been possible. We also thank Peter Traber and Anatole A. Klyosov (Galectin Therapeutics, Newton, Massachusetts 02459) for providing GM-CT-01 and GR-MD-02, and Ravia Salgia for the generous gift of the 3F12 anti-Abl monoclonal antibodies. This study was supported by PHS NIH RO1 CA172040 and CA090321 (to NH).

## Abbreviations

BCP-ALL: B-cell precursor acute lymphoblastic leukemia
CML: chronic myelogenous leukemia
dKO: double knockout
DTSSP: (3,3’-dithiobis(sulfosuccinimidyl propionate))
MEF: mouse embryonic fibroblast
MFI: mean fluorescent intensity
Ph-positive: Philadelphia chromosome positive
PDX: patient-derived xenograft
Ph-negative: Philadelphia chromosome negative
SFK: Src family kinase
WB: Western blot
wt: Wild type

---

**Figure S1.**
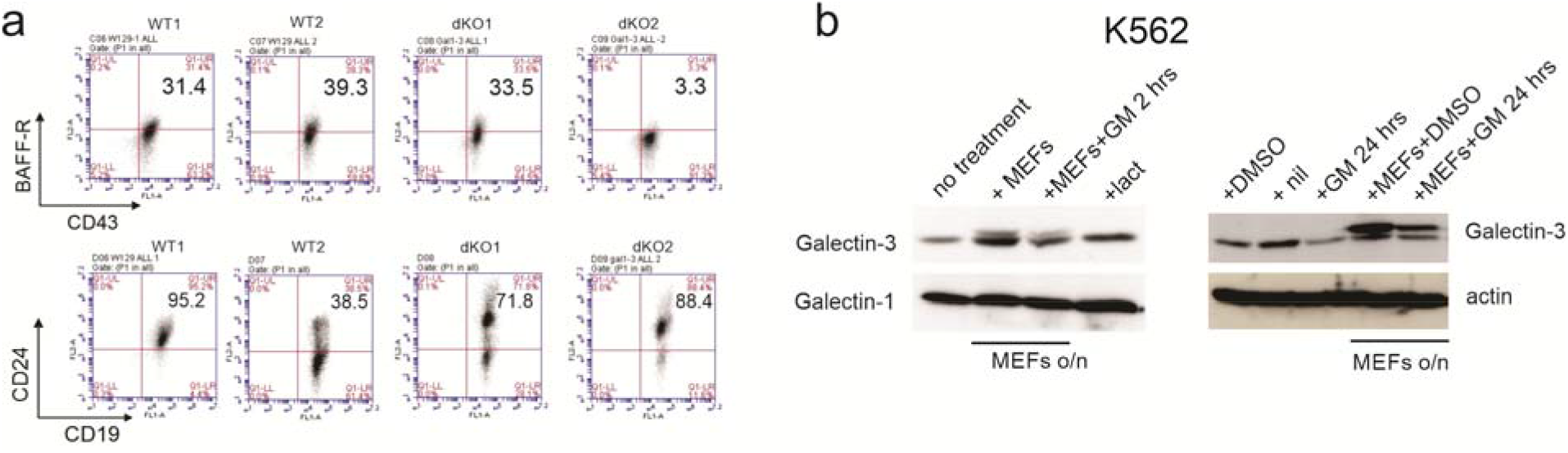
(**a**) FACS analysis of wt1, wt2, dKO1 and dKO2 murine BCP-ALL cells for the indicated cell surface markers. Numbers indicate the percentage cells in the upper right quadrant. (**b**) Western blot analysis of K562 cells grown alone or with irradiated MEFs overnight as indicated. Antibodies are indicated to the left and right of the panels. Actin, loading control. Lane 1 and lane 2, K562; lane 3 K562 incubated with 5 mg/ml GM-CT-01 (GM) for 2 hours; lane 4 K562 incubated with 50 mM lactose (lac) for 2 hours. Lane 5 and lane 8, DMSO-treated K562; Lane 6 K562 exposed to 1 μM nilotinib (nil); lane 7 and lane 9, K562 treated with 2 mg/ml GM-CT-01 for 24 hours.

**Figure S2.**
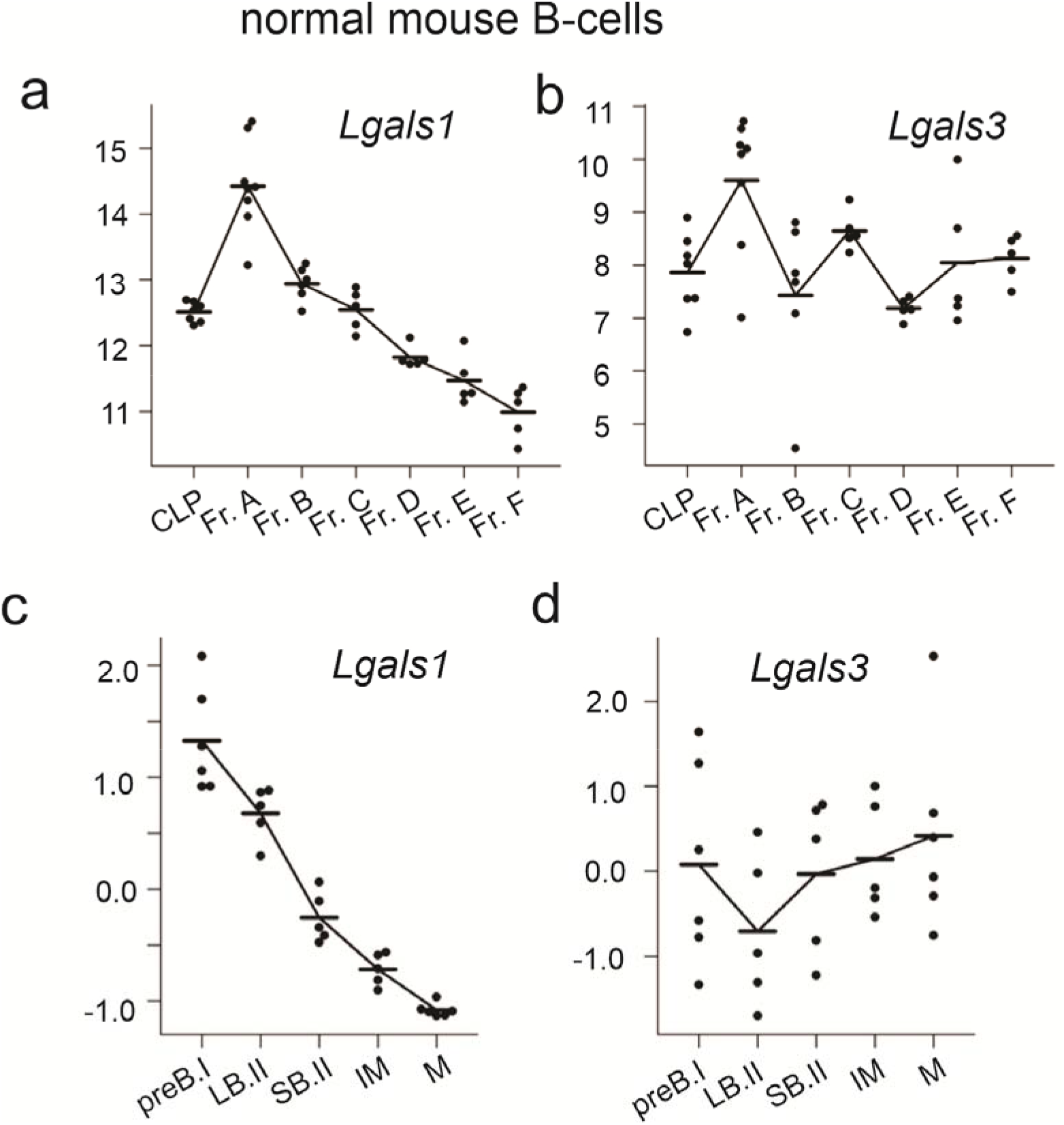
Murine *Lgals1* and *Lgals3* are both expressed during normal B-cell development but in a different pattern. (**a, b**) *Lgals1* and *Lgals3* expression during normal mouse B cell development. Gene expression (GSE38463) of 41 samples from flow-sorted B-cell precursor populations from common lymphoid progenitor (CLP) through to Hardy stage F (83). Values, MFI of individual samples. (**c, d**) Gene expression (GSE13) of 5 stages of normal B-cell development on flow-sorted or *ex vivo* cells including pre-BI, large pre-BII, small pre-BII, immature B and mature B cell stages. Each dot represents an independent replicate. Normalized average difference values as described (84).

**Figure S3.**
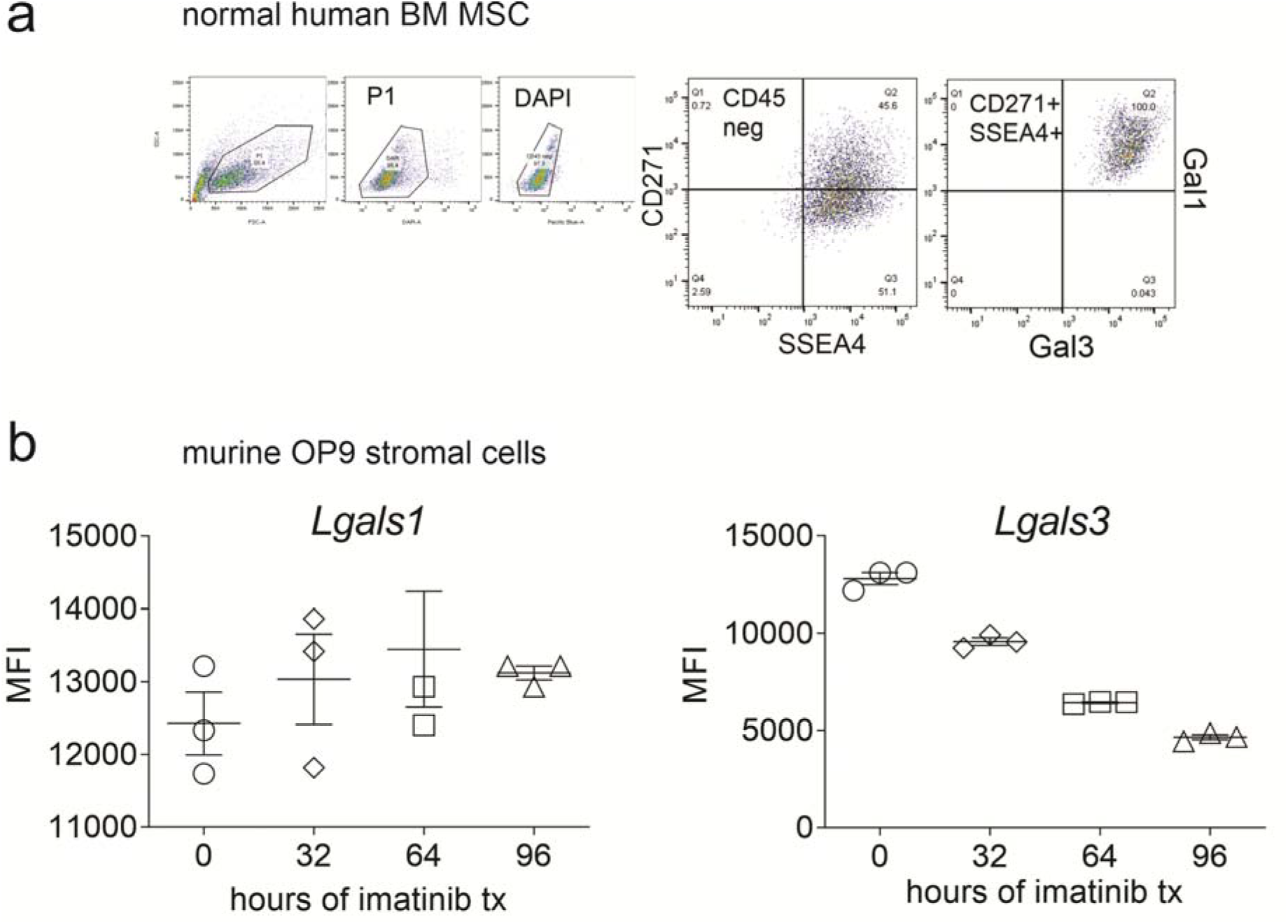
Lgals1 and Lgals3 are both expressed in stromal cells. (**a**) Normal human bone marrow mesenchymal stromal cells described in (24) analyzed by FACS for both Galectin-3 and Galectin-1 expression. One of two samples with similar results. (**b**) Expression (GSE56472) of *Lgals1* and *Lgals3* in OP9 bone marrow stromal cells treated with imatinib for the indicated time. Triplicate samples, Illumina GeneChip (85). Differences between *Lgals1* expression values are not significant, but are significant (p<0.01) at all time points for *Lgals3*. One-way ANOVA, multiple comparisons.

**Figure S4.**
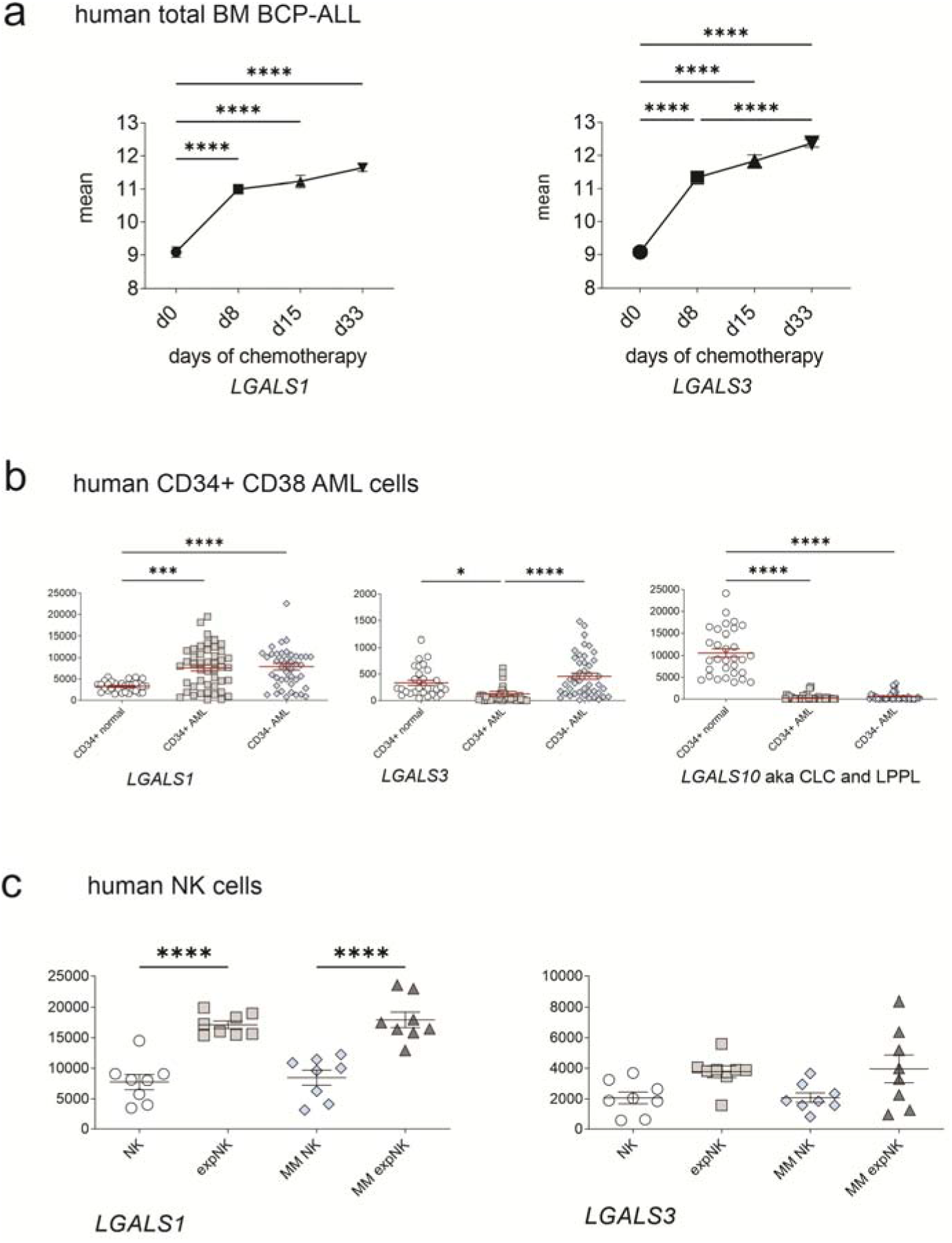
*LGALS1* and *LGALS3* have overlapping expression in normal and abnormal human hematopoietic cells. (**a**) Increasing levels of *LGALS1* in bone marrow of pediatric ALL during chemotherapy treatment. Mean log-transformed normalized GEP values (GSE67684) for the indicated genes on 220 pediatric de novo ALL at diagnosis, day 8, day 15, and day 33 of remission-induction therapy. *LGALS3* also shown in (24). (**b**) *LGALS1* and *LGALS3* expression (GSE30029) compared to previously reported *LGALS10* differential expression (86) in AML cells. Magnetic bead-isolated normal bone marrow CD34+ progenitors (n= 31) compared to flow-sorted AML CD34+ and CD34-mononuclear cells (87). Illumina BeadChip arrays; each symbol, one sample. (**c**) *LGALS1* and *LGALS3* gene expression of human NK cells (GSE27838) from eight healthy donors and multiple myeloma patients, and NK cells from the same two groups after *ex vivo* expansion in the presence of K562-mb15-41BBL cells (88). Log-transformed GEP intensity values. Affymetrix genome arrays. *p=0.0228; ***p<0.001; ****p<0.0001. One-way ANOVA, multiple comparisons, Tukey’s multiple comparison test.

