## Supplementary material for "Cell-Intrinsic and Extrinsic Effects of Galectin-1 and Galectin-3 in B-Cell Precursor Acute Lymphoblastic Leukemia": previous peer review

### Page 1-8 round 1 of review

### Page 9-10 round 2 of review

### **ORIGINAL REVIEW ROUND 1 -cover letter and reviewers #1, #2 and #3 plus our responses**

(1) Cover letter

Dear Editor,

We have uploaded the revised version of our manuscript “Cell-Intrinsic and Extrinsic Effects of Galectin-1 and Galectin-3 in B-Cell Precursor Acute Lymphoblastic Leukemia”.

We note that the contribution of Dr. Francoise Poirier (Morphogenesis, Homeostasis and Pathologies Group Institut Jacques Monod - UMR 7592 CNRS - Université Paris), who provided essential materials, was significant. Before I originally submitted this manuscript I emailed her with a draft but did not receive a response. Since she had not read or approved the manuscript I was going to submit, I could not include her as a co-author, based on what I presume are universal guidelines. With the revision of this manuscript I again tried to establish contact, this time by phoning to Paris. I spoke with Dr. Mireille Viguier, who told me that Dr. Poirier has been in the hospital for a long time, is gravely ill, and will never be able to read the manuscript. Therefore I had no choice but to leave her name in the acknowledgements.

Do you agree that this is the only option or do you see another way to acknowledge her contribution?

We very much appreciate the time and effort of the reviewers who assessed our manuscript and their constructive comments. We have provided our answers below to respond to some of the critiques. We have indicated the modified text in the main body of the manuscript with a different font color.

Sincerely yours,

### ORIGINAL REVIEW ROUND 1

(2) Reviewer #1 and our responses

Major reserves:

**Comment 1.** In the abstract, the authors wrote that galectin-1 and galectin-3 expression patterns overlap in their action and could have potentially common functions… Any references from the past literature? In their work they used double KO for galectin-1 and galectin-3, it is therefore impossible to give an effect to one of them…. I would like an introducing experiment showing the use of Gal-1 KO, Gal-3 KO and double KO vs wt BCP ALL cells

**Response:** There are quite a few cell types that express both Galectin-3 and Galectin-1, at least on a gene expression level. These include for example human natural killer cells (GSE27838), OP9 cells (GSE56472- PMID 25712988) and also equine mesenchymal bone marrow stromal cells (29096716). We have added this information to the discussion section. We agree with the reviewer that a direct comparison of Gal1, Gal3, wt, and Gal1x3 dKO BCP ALL mouse cells would be informative but for practical reasons we were not able to do so. To generate these materials, one needs the mice for isolation of bone marrows. We obtained viably shipped bones from matched wt and double knockout bones from Francoise Poirier and retroviral transduction on these was performed at the same time to reduce biological variation at this step. Unfortunately, Jackson labs (stock 006337) does not maintain Gal1 KO mice as active breeding colonies so this genotype was not available. We note that other studies such as PMID 26250529 on reactive astrocytes also only compared wt and dKOs.

**Comment 2.** Plant compounds GM-CT-01 and GR-MD-02: how are they used? on wt cells? There is nothing about these compounds, and after a look on internet, it appears that a lot of datas were obtained with these compounds. For example, GR-MD-02 targets galectine-3 with higher affinity than galectine-1, and both compounds are already used in clinical trials…. It is really embarrassing to have to wait until the discussion to get interesting datas on these compounds like their affinity

**Response:** We apologize. It is difficult to organize a manuscript in such a way that every reader finds the required information in the desired place. Reviewer #4 remarked “…. The results section contains a lot of discussion moment which must be places in the discussion section. Also, authors mention some methods and approaches in the results and this makes them more difficult to understand”. We had organized our manuscript in this way to provide the reader with information at the locations we thought were specifically relevant. To address the comment of R#1 we have now included information on the compounds in the Introduction section.

**Comment 3.** No statistical methods were described.

**Response:** We have addressed the critiques and comments of all reviewers pertaining to statistical methods and analysis.

Other

**Comment 4:** Line 59: "manuscript in submission"

Figure 1: a: is it significant? LGALS3? LGALS1? No effect of the different treatments in fig 1b right panel.

Line 90: first appearance of nilotinib: what is it? must be explained

Line 96: wt1, wt2, dKO1 and dKO2: should be explained once.

Why fig 2 is placed there? It should be inserted beneath or in paragraph 2.2

Line 116: MS in preparation???

**Response:** Statistics: Please see R#1 C#3 response. Nilotinib as specific inhibitor of the activated Abl kinase is now mentioned. We have also addressed the other points made by this reviewer.

**Comments 5- Fig 3:** I am a bit puzzled by how the results are presented, especially with the dKO cells. In Fig 3a, wt1 and wt2 are a bit different, is it significant? For dKO1 and dKO2 the results are statistically different due to very weak viability. Next on fig 3b, the authors have chosen to use wt2 and dKO2, which are closer than wt1 and dKO1 in term of proliferation… why this couple?

Why on fig 3b, wt2 starts with a 90% viability, and dKO2 cells with only 60 %; could the authors explain ? For fig 3d and e, the authors showed results from wt1 and dKO1, the 2 groups with the largest difference in cell proliferation (fig 3a). To simplify the authors should make a choice between the 2 wt and 2 dKO and explain it, or they must show all the results.

Fig 4: in a, there is a clear difference between the 2 dKO: any explanation?

**Response: Fig. 3a and Fig. 4-** For analysis of the growth rate of the wt and dKO we merged the data for statistical analysis but left the graphs separate. In addition we now have included a second graph of growth to illustrate that, whereas the significant differences between wt and dKO remain, the absolute cell numbers increased upon longer culture. We would like to draw attention to the fact that these dKO cells are not cell lines. The starting material is total mouse bone marrow from matched wild type and double null mutant mice. Populations of precursor B-lineage cells in these bone marrows are transformed with retroviruses that encode an oncogene, Bcr/Abl (Bcr/Abl is also the primary transforming oncogene of responsible for the development of Ph-positive human ALL). Murine precursor B cells that have been successfully transformed by Bcr/Abl are independent of IL7 for their growth. However, dKO1, dKO2, Wt1 and Wt2 initially each still represent batches of polyclonal cells and there is clearly a biological variation. If one would compare different primary human ALLs even from one subtype such as the Ph-positive subclass, one would also measure differences in growth rate in a side-by-side comparison or differences in absolute numbers of migrated cells. Thus WT1/WT2 and dKO1/dKO2 are not identical.

**Fig. 3b.** dKO cells appeared to be more fragile and easily stressed. They grew more slowly and frequently had a somewhat lower initial viability than the WT cells. This was also for example seen in the former fig. 3e where initial viability was around 50%. We did not investigate the mechanism but speculate that loss of endogenous Lgals3 may decrease their overall resilience to stress such as exposure to temperature fluctuations or to 22% O2 used in tissue culture. For example, we saw (unpublished observations) that Lgals3-/- MEFs, compared to WT MEFs, had decreased population doubling levels both at 22% and 6% O2.

**Fig. 3d and fig. 3e**. In the original figure we only showed one set treated with vincristine (wt1 x dKO2) and two with nilotinib (wt2/ wt1 / dKO2 x nil; wt1 x nil + MEFs and dKO1 x nil x MEFs) with the objective of illustrating a variety of drug treatments with the different genotypes. Experiments were not set up with any intent to compare specific combinations, and WT1 and WT2 overall had similar growth. In the revised manuscript we have reorganized the figure which we hope improves the presentation. We now show wt2 / dKO1 treated with vincristine as well as wt1 / dKO2 also treated with vincristine; and the variables MEFs, nilotinib on a wt1 / dKO1 set and on a wt2 / dKO2 set. Although individual experiments have a variation in the absolute cell counts or viability the differences between dKO and WT cells are consistently seen consistent with differences associated with different genotypes.

**Comments 6:** I would like to understand why next, the authors chose to use human TXL2 and US7 cells ? The experiments done with these human cell lines could be done with the murine BCP ALL, and the results with the 2 plant compound could be of interest…

**Response:** As noted in the Materials and Methods of the manuscript, transformation of murine bone marrow cells with Bcr/Abl produces an aggressive pre B leukemia in which the cells are not dependent on stroma for their growth. However, our previous studies with primary PDX-derived human BCP cells show that these strictly depend on the presence of stroma for their growth. Stromal cells synthesize and secrete Galectin-3 and Galectin-1. As the plant compounds are thought to mainly work by inhibition of extracellular (stromal produced) galectin we needed to test their activity in this co-culture system and with human cells.

**Comment 7 -Fig 5:** what about Gal-1 ??? These experiments are focused on Gal-3, and the design of the experiment is really strange… K562 cells express Gal-3, it could be detected intracellularly. Next to investigate the effect of exogenous Gal-3, K562 cells were co-cultured with MEFs… and there is more Gal-3…. Does it mean that this Gal-3 comes from MEFs ? no. Gal-3 could be secreted by K562 cells and MEFs stop this secretion. To clear this trouble, K562 cells are easy to transfect, and siRNA against Gal-3 could be used… the same experiment done with these cells could be of interest.

**Response:** Yes the reviewer is correct. K562 cells produce Gal3 endogenously, as also shown by Yamamoto-Sugitani (PMID 21987825). K562 cells also contain Galectin-1. In the revised manuscript in Supplementary Figure 1 panel b we have included a Western blot for Galectin-1 of K562 cells treated with GM-CT-01. However, the signal for total Galectin-1 (we can not distinguish murine and human Galectin-1 as they have the same size) is similar in all lanes. The co-culture with stroma increases the Galectin-3 in K562, but this is made by the stroma: in our experiments, the extra Gal3 is from the MEFs- we have previously published that stromal cells secrete Galectin-3 and that it is taken up by leukemia cells. Our published and unpublished data show that stromal produced Galectin-3 is protective for leukemia cells.

**Comment 8:** Fig 6a: the effect of GM-CT-01 seems tiny… I do not understand why the authors used GM instead GR? GR binds Gal-3 with higher affinity and can target extracellular Galectin…

Fig 6b: lactose has no effect, that is surprising for the well-known inhibitor of Galectins… could the authors discuss this point ?

A 2d band is viewable in presence of MEFs, and the authors wrote this is due to the murine gal-3. A western blot with (purified) human and murine gal-3 will help…

Discussion: line 290: at least ! I had to almost finish the reading of the MS to get important data on the 2 plant compounds…

**Response:** Fig 6a- we agree that the effect of GM-CT-01 is small. Such compounds appear to be mainly cytostatic in tissue culture. Regarding the choice between GM-CT-01 and GR-GM, GM-CT-01 was originally known as Davanat, was the first compound used to treat human patients and was the first of the two compounds the company made available to us. Interestingly, based on published in vitro data, the Kd of both GM-CT-01 and GR-GM is similar for both Galectin-1 and Galectin-3 which could lead to the expectation that they have similar potencies in tissue culture. Clearly, in a tissue culture setting, the strength of their effect is not the same. This can be caused by for example differences in degradation.

Fig. 6b. We agree with the reviewer: lactose is a well-known, albeit aspecific inhibitor of Galectin-3 that is needed at high concentrations. However it difficult to predict what its expected effect would be under the experimental conditions used by us:

Lactose as Galectin-3 inhibitor reduces binding of Galectin-3 to its target glycoproteins. As such interactions take place extracellularly, lactose is, for example, used to cause the displacement of extracellular Galectin-3 bound to the surface of cells as measured using flow cytometry. We have used lactose for this purpose in combination with exogenously added recombinant Galectin-3 and it very clearly displaces Galectin-3 from the cell surface.

If lactose treatment reduces Galectin-3 levels seen on a Western blot of a total lysate will depend on where that Galectin-3 was located. We note that K562 constitutively synthesizes its own Galectin-3 endogenously and thus one can assume that a varying percentage of that is exported and presented on the surface. The extracellular Galectin-3 is expected to be knocked off the cell surface by lactose, and will be gone when the cells are washed and harvested for the Western blot. One would not expect intracellular Galectin-3 to be affected by lactose. Thus the endogenous intracellular Galectin-3 constitutively expressed in K562 should remain the same with or without lactose. Based on molecular weight (see below) we can identify extracellular Galectin-3 made by the mouse stromal cells. We do not know how much of it or how rapidly it is endocytosed. However, our results show that GM-CT-01 mainly reduced the mouse Galectin-3 band, consistent with the idea that it displaces cell surface Galectin-3. In Fig. 6b, a small reduction of the Galectin-3 signal was seen in the lactose-treated K562 but the effect on the mouse Galectin-3 was indeed very small. We have included other similar analysis in supplementary Figure S1 (panel b). We have included this discussion in the Discussion section of the manuscript.

As mentioned above, the second Galectin-3 band seen in this Western blot is definitively mouse Galectin-3. This is because, interestingly, human and mouse Galectin-3 are *different in molecular weight*: human Gal3 (UniProt P17931) is 250 aa whereas mouse Gal3 (UniProt 16110) is 264 aa. The human protein lacks residues at around amino acid residue 67 in the repeat domain. Mouse and hamster Galectin-3 have the same molecular weight. We have added this information to the manuscript and show this in the supplemental Figure 1c. To the best of our knowledge there have been no studies that examine if the difference results in functional differences between human and mouse/hamster Galectin-3.

### ORIGINAL REVIEW ROUND 1

(3) Reviewer R#2 and our responses

**Comment Introduction:** The term “Ph-chromosome” “Ph-positive and Ph-negative leukemia” should be described.

**Reply:** We have addressed this in the revised manuscript.

**Comment Results:** Fig 1a, what was the rationale for analyzing Ph-positive ALL in two gene arrays? In addition, the expression of galectin-1 and galectin-3 in Ph-positive ALL was shown. Although the author claimed that many samples showing a positive correlation between expression of these two genes. However, no statistical analysis was not performed. How about the expression of galectin-1 and galectin-3 in other types of ALL?

**Reply:** Based on DNA sequencing and the identification of driver and other oncogenic mutations, the amount of different subcategories into which human BCPALL can be subdivided has expanded in the past 25 years, and some of these genetic lesions also have prognostic value. This poses both a problem and an advantage for investigators: which, if any subcategory to choose for study? For example, Ph-positive leukemia, and Ph-like ALL are characterized by the constitutive activation of signal transduction pathways whereas other BCP ALLs are driven by abnormal transcription factors. Because the mouse model used by us and others is driven by Bcr/Abl (the molecular culprit in Ph-positive leukemias) we compared Ph-positive human ALLs, and also included the CML cell line k562 in our protein analysis because this type of leukemia, although from the myeloid lineage, is also caused by Bcr/Abl. In the revised manuscript we have now also included the MLL and the Etv6/Runx1 (aka tel-AML) subcategories which involve transcription factors.

Statistics: Pearson correlation analysis has now been performed for these data and its statistical significance analyzed in the revised manuscript (Figure 1a, b).

**Comment 2 Results:** Fig. 1b, was the results analyzed the dataset from reference 21 and published by the authors’ lab? Please upload it to GEO. In addition, the abbreviation in Fig 1b should be described in figure legend.

**Reply:** The dataset in Fig. 1b was generated in a collaboration between our lab and the Muschen lab. Our lab generated the samples and they did the gene expression analysis. They also used the data in a publication, which we cited in our original manuscript. However, in response to the reviewer’s question I went to GEO to verify that the data had been deposited and discovered that they had not in fact been deposited there. Fortunately, we found the original datasets and submitted them to GEO. They are now publicly available. We thank the reviewer, as we put quite some energy into generating the samples and agree that such data should be publicly available.

**Comment 3.** The protein expression of galectin-3 must be shown in Fig 2b.

**Reply:** We have now included this in Fig 2b; on the original Western blot, apart from the positive control, not much was visible in any of the mouse BCP ALL samples. Human BCP ALL cells make little Galectin-3 endogenously under steady state conditions in tissue culture and it appears the mouse BCP ALL cells are similar. The human cells contain Gal3 secreted by the OP9 cells but, as described in the manuscript, the default culture for the mouse cells is without stromal support.

**Comment 4.**Line 110, “However, levels of p-Src were reduced in the dKO cells.” However, the level of p-Src in dKO should be increased in Fig 2b?

**Reply:** We sincerely thank the reviewer for catching this rather embarrassing mistake! Levels of phosphorylated Y416 on pSrc /Src family kinases were indeed increased. As we have not noted this in our Gal3 KO cells we speculate the increased pSrc may relate to the regulation of transmembrane tyrosine phosphatase CD45 by Galectin-1. Dephosphorylation of phosphotyrosine Y416 of Src de-activates its kinase activity. Thus, deletion of Galectin-1 could reduce CD45 activity on this residue and the increased pSrc /Src family kinase pY416 (for example on Lyn or Lck kinases) could be explained by this mechanism. Reviewer #3 remarked that this should be discussed. This is quite speculative, but we have incorporated this in the discussion.

**Comment 5.** Fig 3, the symbol **p<0.01 ***p<0.001 ****p<0.0001 should be indicated the difference between specific control and experimental group.

**Reply:** We have improved the figure legend to indicate which comparisons were done**.**

**Comment 6.** Besides inhibitors, siRNA experiment is essential to demonstrate the endogenous effect of Galectin in human ALL.

**Reply:** Although we agree with the reviewer that siRNA could be an alternative or complementary approach, there are significant technical hurdles that preclude this in the cell type that we use, which are primary, PDX-derived human patient samples. We are not able to introduce nucleic acids into these cells with an appreciable efficiency, and even lentiviral transduction at best delivers 4% transduced cells. In our hands, transfection, nucleofection and electroporation all have efficiencies that are even lower. Thus, it is not feasible to generate galectin-1 x galectin-3 dKO human ALL cells. We are currently attempting to generate knockout cells using the CAS/CRISPR system but are, also in this case, faced with the low efficiency of the introduction of nucleic acids into this cell type.

**Comment 7.** Exogenous effect of galectin-3 was examined by coculture system. Because no evidences supported high galectin-3 concentration in OP-9 coculture system, antibody neutralizing experiment or recombinant galectin-3 treatment should be performed in order to confirm the exogenous effect.

**Reply:** This comment was not clear to us. We have previously examined and published that OP9 cells synthesize and secrete abundant Galectin-3 (Fei et al, 2015), and we showed the effect of stimulation of human BCP ALL cells with (recombinant) exogenously added Galectin-3.

**Comment 8.** The effect of galectin-3 was investigated in BCP ALL US7 (Ph-negative) and BLQ1 (Ph-positive). Similar signaling.

**Reply:** It was not clear to us if the reviewer expects a reply to these comments.

**Comment 9.** Why did the expression of galectin-1 and galectin-3 in Ph-positive was investigated by meta-analysis? Why was K562 used in some experiment? The rationale should be clarified.

**Reply:** Please see the response to R#2 who also asked about the choice of cells. We have added a clarification to the revised manuscript.

**Comment Materials and Methods**

1. The procedure of murine MEF isolation is missing.

2. No statistical methods were described.

**Reply:** 1. Isolation of such cells is a relatively standard procedure. However, we have now added a brief description of the methods used by us. 2. Statistics: Please see R#1 C#3 response.

### ORIGINAL REVIEW ROUND 1

(4) Reviewer #3 and our responses

**Comment** Line 59: typo - 'manuscript submitted'

Figure 1 is there significance for a? What statistical test was used for b, and is there a 'mock' treatment for nilotinib (eg DMSO)? How much nilotinib was given?

**Reply:** Yes, we only have one manuscript on our studies on Galectin-1 and that is currently under peer review. Statistics: Please see R#1 C#3 response.

Yes control for nilotinib always was DMSO. We have also expanded the text in the Materials and Methods section.

**Comment** **Figure 2a.** If possible a third mouse would be good as CD43/B220 for dK02 appears slightly different. Also, can reviewers see BAFFR and CD24?

**Reply:** Please see response to R#1 comment 1 and comment 5. We have included the FACS plots for the BAFFR and CD24 for the reviewer. We did not see a genotype-correlated difference in these cell surface proteins so we assume this is genotype-independent biological variation of these primary samples.

**Comment** **Figure 2b.** We need to see absence of galectin-3. Can you include the galectin-3 blot mentioned in the text? It doesn't matter if it is untidy. Quantification using densitometry, normalized to GAPDH, would be very useful for the reader. Need a comment on p-Src in the discussion. It appears to be a clear result and ideally would be commented upon.

**Reply:** The issue of the Galectin-3 WB was also raised by R#2 (comment 3)- please see that response. We have quantified the differences by densitometry of the pSrc/Src ratio. This is now included in the figure. The issue of p-Src was also raised by R#2 (comment 4). Please refer to the response.

**Comment Figure 3.** How was viability measured? What statistical tests are used? **Comment** **Figure 4**. What statistical tests are used? Did the compounds have any impact on cell health?

**Reply:** Viability: we have included more description in the materials and methods section. Statistics: We have re-analyzed all the data for statistical significance and included information on the analysis in the Materials and Methods.

As measured by viability, the compounds as mono-treatment had no effect on the health of the cells. This is consistent with studies in mice and in people where the compounds do not show toxicity.

**Comment** **Figure 5a**. Do you have the missing part of the Western - top right - total lysate?

**Reply:** We have included the missing part. That particular anti-c-Abl antibody produced a weaker signal than the 3F12 antibody, and that WB is an overexposure. We have noted the differences in exposure levels of the panels in the figure legend.

**Comment** **Figure 5c**. Can you improve labeling and explanation in legend - for lanes 5 and 6? Suggest include 'respectively' after text explanation.

**Reply:** We agree with the reviewer- these results were presented in a confusing way. We have now enlarged the relevant parts of Figure 5c, and provided a more detailed figure legend that makes use of the numbers of the lanes.

**Comment** **Figure 6.** Just a quick comment to reviewers about the concentration of GM-CT-01 - does it need to be so high? Densitometry for these blots could be useful for the reader.

**Reply:** These compounds are lower molecular weight derivatives of complex, plant-derived carbohydrates. Nonetheless, based on their still substantial size (50-60 kDa) they will need some type of active transport to enter cells. We assume that they mostly remain extracellular and so their main mechanism of action may be displacement of Galectin-3 and Galectin-1 binding to cell surface glycoproteins. The affinity for Galectin-3 and -1 binding to such compounds may actually not be so high, or be lower than the affinity of Galectin binding to the natural ligands, and this may account for the high concentrations needed to obtain an effect.

**Comment** **Figure 7**. Can you indicate if a DMSO control was used for the nilotinib?

**Reply:** Yes, DMSO was the control. We have included this in the figure legend.

(5) Reviewer #4 and our responses	Comments and Suggestions for Authors

**Comment 1.** The manuscript lacks description of statistical evaluation. There is no information how the differences between values were measured, what tests were performed and so on. Without statistics all the results are useless.

**Reply:** We agree. Statistics: A paragraph describing the statistical analysis methods used has been added to the Materials and Methods section. Also, all graphs have been re-analyzed. Results are also described in the figure legends.

**Comment** **2**. Author did not mention how the cell viability was measured. Also, Western blot analysis methods are not described. The murine MEF isolation methods are not described at all. In other words, Materials and Methods section must be re-worked and all methods used should be described in detail.

**Reply:** We tried to follow the Instructions to Authors for the IJMS as described online: “the methods should be described with sufficient detail to allow others to replicate and build on published results. New methods and protocols should be described in detail while well-established methods can be briefly described and appropriately cited.” Because cell viability counting, Western blot analysis and MEF isolation are regarded as routine, we did not provide much detail. To address this reviewer’s comment, we have now included a short section providing more details of the methods used for these experiments.

**Comment** **3.** In figures the symbols **p<0.01 ***p<0.001 ****p<0.0001 should be indicated the difference between specific control and experimental group.

**Reply:** Detailed annotation of p-values using ** is specified in the figure legend.

**Comment** **4**. The whole manuscript is poorly organized. The results section contains a lot of discussion moment which must be places in the discussion section. Also, authors mention some methods and approaches in the results and this makes them more difficult to understand.

**Reply:** We apologize. How to optimally organize a manuscript is difficult. If no explanation is given for a certain approach in the Results section for example, it may be difficult to follow the logic of why an experiment was done. Some reviewers and authors make a strict separation of the different sections whereas others try to provide some information at the place where it is needed. We have attempted to find a balance.

### ORIGINAL REVIEW ROUND 2 – letter from the guest editor and our response

### (1) Letter from guest editor Dear Prof. xx

### Thanks for your comment that was fully considered by the Editor. I can easily understand your disappointment regarding the Editor’s decision to reject your manuscript. I personally experienced several times this stressing condition. Your work is potentially interesting and almost ready for publication in my point of view, but unfortunately it lacks of some critical experiments that were suggested by one of the reviewers during the first round revision. These experiments were not considered at all in your revised manuscript…. The problem is simple: You have got comments about your paper by a panel of four reviewers (this is an opportunity for you and not a damage.....), three of which were positive; the first one, which “unfortunately” is the expert scientist in the field, asked for specific experiments in order to reinforce the paper quality and the authors’ conclusions…. His request sounds pertinent. This does not mean that all the suggested experiments have to be necessarily performed, but you should reply to these comments adequately… ! (I will give you my specific opinion about this below….) A point by point reply to the reviewers’ comments is required with the aim to resolve the criticisms. In order to be accepted for publication the revised manuscript should satisfy all the pertinent questions / comments posed by the reviewers. This is the rule for any scientific journal….You wrote 8 page-response to the reviewer 1, without addressing such a critical points! Your reply to the major comment 1 is scientifically unacceptable. You are presenting data with double KO cells but you cannot set-up an in vitro model with a single KO?. The reviewer cannot accept this response and the editor of a scientific journal cannot fly on this specific issue…. Below is reported the integral comment 7 by reviewer 1 and your specific reply….. Just to give you a further example: Comment 7 -Fig 5: what about Gal-1 ??? These experiments are focused on Gal-3, and the design of the experiment is really strange… K562 cells express Gal-3, it could be detected intracellularly. Next to investigate the effect of exogenous Gal-3, K562 cells were co-cultured with MEFs… and there is more Gal-3…. Does it mean that this Gal-3 comes from MEFs ? no. Gal-3 could be secreted by K562 cells and MEFs stop this secretion. To clear this trouble, K562 cells are easy to transfect, and siRNA against Gal-3 could be used… the same experiment done with these cells could be of interest.

### Response: Yes the reviewer is correct. K562 cells produce Gal3 endogenously, as also shown by Yamamoto-Sugitani (PMID 21987825). K562 cells also contain Galectin-1. In the revised manuscript in Supplementary Figure 1 panel b we have included a Western blot for Galectin-1 of K562 cells treated with GM-CT-01. However, the signal for total Galectin-1 (we can not distinguish murine and human Galectin-1 as they have the same size) is similar in all lanes. The co-culture with stroma increases the Galectin-3 in K562, but this is made by the stroma: in our experiments, the extra Gal3 is from the MEFs- we have previously published that stromal cells secrete Galectin-3 and that it is taken up by leukemia cells. Our published and unpublished data show that stromal produced Galectin-3 is protective for leukemia cells.

### Do you feel that comment 7 has been fully addressed ?…. And what about the suggested experiments to be performed with gal-3 and gal-1 specific mRNA interference in K562 cell line? These experiments do not require one year work ! I can understand that the request to make galectin-3 and Galectin-1 specific KO mice is a little bit hard to digest considering the hard work necessary to create and characterize these in vivo models (specific gal-3 KO mice are however described in the literature and are probably available) but please, to get specific K562 KO cell lines for gal-3 and Gal-1 to be used in selected experiments as suggested, does not require a big effort! With these interfered K562 cell lines some in vitro experiments can be improved according to the reviewer’s suggestions and the manuscript can be finally accepted…. It is pacific that pertinent criticisms and suggestions posed by a competent reviewer cannot be fully skipped !. Just to stress again this concept: Your statement ……. “The co-culture with stroma increases the Galectin-3 in K562, but this is made by the stroma”……:This represent the authors’ opinion not a scientific evidence !!! By using the aforementioned gal-3 KO K562 cell line you can easily resolve this issue….. Concluding, this is the reason why your potentially interesting manuscript has been rejected with the possibility of resubmission. If Editors invite competent reviewers to express an opinion about a scientific work, we should accept their comments and when the resolution of specific problems is possible with simple experiments the author should perform such a work in order to address each specific comment or eventually the authors can reply to the comment in a scientific way. Sincerely Yours Guest Editor IJMS

(2) Our response to letter from the guest editor

It is the opinion of the guest editor that for acceptance for publication to this journal all comments from each of four different reviewer need to be addressed, even though it concerns doing extensive additional bench work experiments. We note that three of the four reviewers found the manuscript acceptable for publication but that one reviewer, according to the guest editor, insisted on more bench work and experiments.

The authors do not agree with this opinion for a number of reasons. (a) We do not agree with the opinion that knocking out Galectin-1 or Galectin-3 or both in K562 cells is of value to the main conclusions of current study. Our main focus is on drug treatment of B-cell precursor acute lymphoblastic leukemia, and *not* the erythroid/myeloid leukemia represented by the K562 cell line. Myeloid and lymphoid leukemias are very different types of malignancy and if one is interested in drug resistance in BCP-ALL, K562 is just not suitable as a model. Also siRNA as suggested by the reviewer gives transient knockdown, and this is not useful if one wants to study long-term development of drug resistance.

That being said, we did extensive experiments to attempt to knock out Galectin-1 and Galectin-3 separately in patient-derived BCP-ALL cells. These are complicated experiments: the cells need to be co-cultured with stromal cells to keep them alive, but of course the stromal cells produce and secrete both Galectin-1 and Galectin-3. This makes screening to monitor successful knockout using FACS for cell surface expression very complicated since no commercial antibodies distinguish mouse from human Galectin-1 or Galectin-3. Overall, we spent a considerable amount of time to try and knock out Galectin-3 in patient-derived BCP-ALL cells using Cas9/CRISPR, without succeeding. We also spent considerable time and effort over a number of years to try to knock out Galectin-1 in BCP-ALL cells, first using lentiviral transduction, then Cas9/CRISPR. We did not obtain single knockout cells for either, and did not attempt to make a double knockout.

(b) We do not fully understand the comments of the editor about our response #7 and why that does not satisfy reviewer #1. If we assume that the sole purpose to knock out Galectin-1 or Galectin-3 or both in K562 would be biochemical and not functional (see point 1 above), it appears that the main point of using K562 would be whether or not the extra band of Galectin-3 seen on the Western blot upon co-culture of K562 with MEFs is murine Galectin-3. The guest editor wrote “does it mean that this Gal-3 comes from MEFs ? no. Gal-3 could be secreted by K562 cells and MEFs stop this secretion”. We do not understand this thought. If K562 cultured alone gives a band on a Western blot and when co-cultured with MEFs a second band appears, given the fact that Galectin-3 in mouse and human differ in molecular weight, the conclusion seems logical that the new second band is mouse Galectin-3. We note that we recently published a peer-reviewed paper <https://pubmed.ncbi.nlm.nih.gov/34830047/> [also see supplementary data associated with that publication] in which we report, among others, the knockout of Galectin-3 in mouse OP9 stromal cells and very clearly make the point that OP9 cells are a major source of Galectin-3 for BCP-ALL cells; also one can see the difference in molecular weight on a Western blot between mouse and human Galectin-3. In our opinion this also addresses the final comment of R#1.

We have revised the discussion of the current manuscript extensively to incorporate the latest data relevant to Galectin-1 and Galectin-3 overlap, and have added new supplementary figures S2-S4.
